# Monocyte-amplified transcriptional signatures of human diseases

**DOI:** 10.64898/2026.07.04.736372

**Authors:** Mario L. Arrieta-Ortiz, Wei-Ju Wu, Nitin S. Baliga

**Affiliations:** Institute for Systems Biology, 401 Terry Ave N, Seattle, Washington, USA

## Abstract

Blood-based biomarkers discovered by machine learning often lack disease specificity and cross-population robustness for clinical applications. We describe a biomarker discovery strategy that exploits monocytes as circulating sentinels to amplify disease-perturbed signals in blood. This strategy leverages monocyteMINER, a mechanistic transcriptional regulatory network inferred from monocyte transcriptomes of 1,202 healthy individuals. As proof-of-concept, we uncovered a 31-gene atherosclerosis-perturbed network that underpins disease etiology, identifying diagnostic signatures for coronary artery disease (ARAP2, P2RY14, FKBP15) and acute myocardial infarction (SERPINA1, ASGR2). For tuberculosis (TB), monocyteMINER uncovered a 5-gene signature (MAS_TB_META5: ANKRD22, AIM2, VAMP5, GBP5, TGM2) from just 438 samples. MAS_TB_META5 outperformed 77 existing signatures across 18 cohorts (>4,400 patients, 12 countries), achieving WHO target product profile for high-sensitivity screening (including in advanced HIV patients), and predicting TB progression up to 5 years before diagnosis. Thus, our findings show that monocyteMINER offers a generalizable platform for discovering clinically actionable biomarkers for diverse diseases.

## INTRODUCTION

Despite the distinct biological mechanisms underlying each pathological condition, including substantial variability in affected organ and tissue localization, most biomarkers discovered through brute force approaches are not disease-specific^1^. For diagnostic biomarkers to achieve disease specificity, it has been proposed that they must originate from abnormal pathology-perturbed tissues or infectious agents and demonstrate direct association with disease etiology and progression^2^. For tissue-derived biomarkers to be detectable in blood, the disease must have progressed enough to raise analyte concentrations above the detection threshold of analytical technologies^3,4^. These constraints make it extremely challenging to discover biomarkers that are detectable in early asymptomatic stages of most diseases, when they are preventable or curable^5,6^. In this regard, blood diagnostics based on transcriptional biomarkers of disease are becoming essential tools in healthcare, facilitating early disease detection, personalized treatment plans, and improved patient management for diverse medical conditions, including infectious diseases, autoimmune disorders, and chronic illnesses^7–11^. However, a generalizable approach for the discovery of biomarkers that are effective in detecting a wide-range of diseases with high sensitivity and specificity, and across diverse patient populations has remained elusive.

Transcriptional biomarkers for a disease are typically discovered through machine learning^12^ (ML) on a large dataset of curated transcriptomes from individuals that are healthy and those that have a confirmed diagnosis, followed by validation of performance on independent patient cohort datasets^3^. Even when such datasets are available, due to similarities of transcriptional response across disparate diseases, and variability in disease dynamics, such as due to host and pathogen genetic backgrounds, there are significant challenges in finding clinically-relevant biomarkers that are both sensitive and specific in detecting a particular disease, while also being generalizable across patient populations and disease contexts^12^.

We sought to address challenges in biomarker discovery by developing an approach that is generalizable but also founded in causal and mechanistic underpinnings of disease biology^12^. There is growing evidence that monocytes are a primary source of transcriptional signatures for many diseases, including COVID-19^13^, TB^14^, Parkinson’s^15^, Alzheimer’s^16^ and cancers^17^. One possible explanation could be that by circulating through blood vessels, and internalizing information through interaction with other cell types and their secreted signals, monocytes serve as an extravascular surveillance system for continuous monitoring of the health status of major organs in the human body^18–20^. There is also substantial evidence that the monocyte population expands during disease, e.g., in TB^21^, Lyme^22^, and cancer^23^. Thus, we hypothesized that decoding the information captured within the transcriptome states of monocytes would denoise transcriptome profiles from whole blood to facilitate the discovery of highly sensitive molecular biomarkers that are causally and mechanistically associated with the biology of a specific disease, and therefore likely to have robust performance across populations and disease settings.

Predictive transcriptional biomarkers that are generalizable across patient populations and diverse disease contexts is especially urgent for accurate diagnosis of pulmonary tuberculosis (TB), which was responsible for ∼1.25 M deaths in 2023^24^. The World Health Organization (WHO) has included culture-free diagnostics that can be easily deployed at point-of-care in remote locations as a core component of its strategy to control and reduce transmission^25^ and end the TB pandemic^26^. Current TB diagnostics rely on bacteriological culture of *Mycobacterium tuberculosis*^27^ (Mtb), sputum smear microscopy^25,27^, assaying immune response to TB antigens (e.g., TB skin tests (TST), Interferon Gamma Release Assays (IGRAs)^27^), which have high false positive rate due to long-term immunologic memory of a prior or a latent TB infection (LTBI)^28^, and detection of Mtb DNA sequences (using the Xpert MTB/RIF molecular assay on sputum)^27,29,30^. Importantly, none of these assays are optimal for diagnosing extrapulmonary TB or pulmonary TB in populations that do not generate sputum (e.g., children^25^). Hence, there is a need for better methods to discover transcriptional biomarkers of TB that have robust performance across diverse patient populations and disease contexts.

Here we present monocyteMINER, a mechanistic systems biology framework that uses a monocyte cell type-specific transcriptional regulatory network to decode and amplify disease-perturbed signals in whole blood. We validate the approach in atherosclerosis, then apply it to TB to discover MAS_TB_META5, a 5-gene signature that outperforms all 77 existing TB signatures in multipurpose utility across 18 independent cohorts, achieving WHO target product profile criteria for screening, detecting TB co-infection in advanced HIV patients, and prediction of TB progression up to 5 years before diagnosis.

## RESULTS

### Framework to discover monocyte amplified signatures of human disease

We built monocyteMINER by applying the MINER algorithm^31^ to monocyte transcriptomes from 1,202 ethnically diverse “healthy” individuals (MESA cohort^32^), yielding a transcriptional regulatory network of 10,567 genes in 5,682 regulons regulated by 529 transcription factors (**Fig. 1A**). As proof-of-concept, applying monocyteMINER to CAC-score-stratified monocyte transcriptomes revealed a 31-gene atherosclerosis-perturbed network centered on HSF1-driven IL-6 inflammation, from which we derived blood-based diagnostic signatures for coronary artery disease (ARAP2, P2RY14, FKBP15; AUROC ≥0.714) and acute myocardial infarction (SERPINA1, ASGR2; AUROC ≥0.814) in independent cohorts^33,34^ (**Fig. 1B-G** and **Table S1**; full results in **Extended Data**). These findings demonstrated that monocyteMINER can decode mechanistically coherent biomarkers even from imperfect training surrogates and small cohorts.

**Figure 1.**
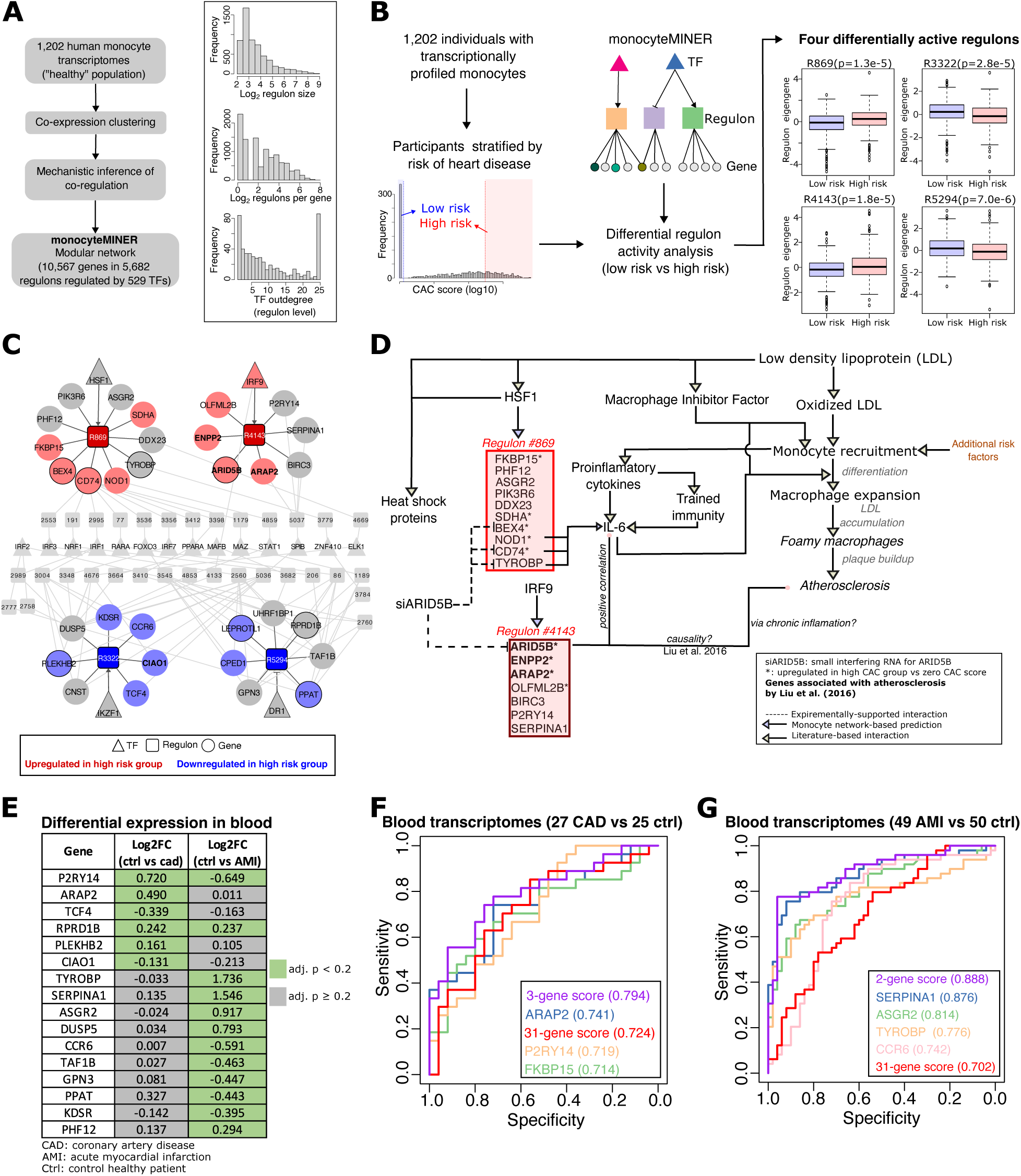
Overview of a monocyte centric framework for discovery of transcriptional signatures for human diseases. A) Pipeline for inferring a modular model for the transcriptional network of human monocytes (i.e., monocyteMINER) using the MINER algorithm^31^. General statistics of the resulting network model are shown as histograms. The monocyte transcriptional compendium used for inferring the monocyteMINER model was sourced from the Multi-Ethnic Study of Atherosclerosis (MESA) by Liu et al^32^. B) General strategy for identifying disease-dysregulated subnetworks in the monocyteMINER model. To identify heart disease risk-associated regulons, the MINER-computed eigengene of each regulon was compared using a t-test for the individuals with CAC scores = 0 (low risk group; n=336) and CAC score > 300 (high risk group; n=293). Only adjusted p-values < 0.05 were considered significant. C) The four regulons identified as differentially active with the workflow in panel B are shown as the nodes with larger size and black edges. TFs associated with these four regulons are also displayed. The sign of the interaction between the TFs and their regulons was defined as the sign of the Pearson correlation between TF expression profile across the 1,202 samples in the MESA transcriptional compendium and the eigengenes computed by MINER for each regulon. Genes associated to atherosclerosis in Liu et al^32^ are shown in bold font. Genes whose expression was affected by a small interfering RNA targeting ARID5B are shown as nodes with a black border. TFs that putatively co-regulate genes from multiple differentially active regulons (and the regulons mediating those connections but not identified as differentially active) are also shown in grey color. Genes for which Wilcoxon rank sum test comparing expression in high-risk vs low-risk groups had p-value ≤ 0.01 are color-coded as indicated in figure key. D) The monocyteMINER recapitulated previously documented association between ARID5B and atherosclerosis via regulon # 4143, while uncovering new molecular interactions and genes (i.e., regulon # 869) that may contribute to the development of atherosclerosis. E) Subset of genes from heart disease risk-perturbed subnetworks shown in panel D that were differentially expressed (i.e., adjusted p-value < 0.2) in blood transcriptomes of individuals in a coronary artery disease cohort (CAD; GEO accession ID: GSE202625^33^) or in an acute myocardial infarction cohort (AMI; GEO accession ID: GSE66360^34^). F) Performance shown as a receiver operating characteristic (ROC) for popular disease score format, defined as the difference between the geometric mean of genes in regulons # 869 and # 4143 and the geometric mean of genes in regulons # 3322 and # 5294 in log2(CPM+1) units, to distinguish patients with CAD from healthy individuals. The ROC curves for three genes within the atherosclerosis network with individual area under ROC curve > 0.7 and their combination (i.e., the 3-gene score) are also displayed. G) Same as panel F, performance of the implemented disease score (and single genes with AUROC > 0.7 and the combination of the two genes with AUROC > 0.8) to distinguish AMI and control cases is displayed. Corresponding AUROC values are shown in parentheses.

### Discovery of monocyte-amplified gene signatures for pulmonary tuberculosis

In order to discover a gene signature that could detect active TB (aTB) with high sensitivity and specificity, while differentiating individuals that are healthy (TB-), have LTBI, or other respiratory diseases (OD), we used monocyteMINER to analyze compendia of transcriptomes from two independent cohorts—one with 11 patients (7 aTB and 4 TB-)^35^ and a second cohort of 181 patients (54 aTB and 127 OD)^36^ (**Fig. 2A**). We describe below how through each of these comparisons we discovered two three-gene signatures that were combined to generate a multipurpose five gene signature. We demonstrate that the five-gene signature has robust performance in detecting aTB across cohorts from different geographic locations as well as in advanced HIV patients, and also in predicting progression in high-risk populations, making it potentially useful across varied contexts, including community-based active case finding.

**Figure 2.**
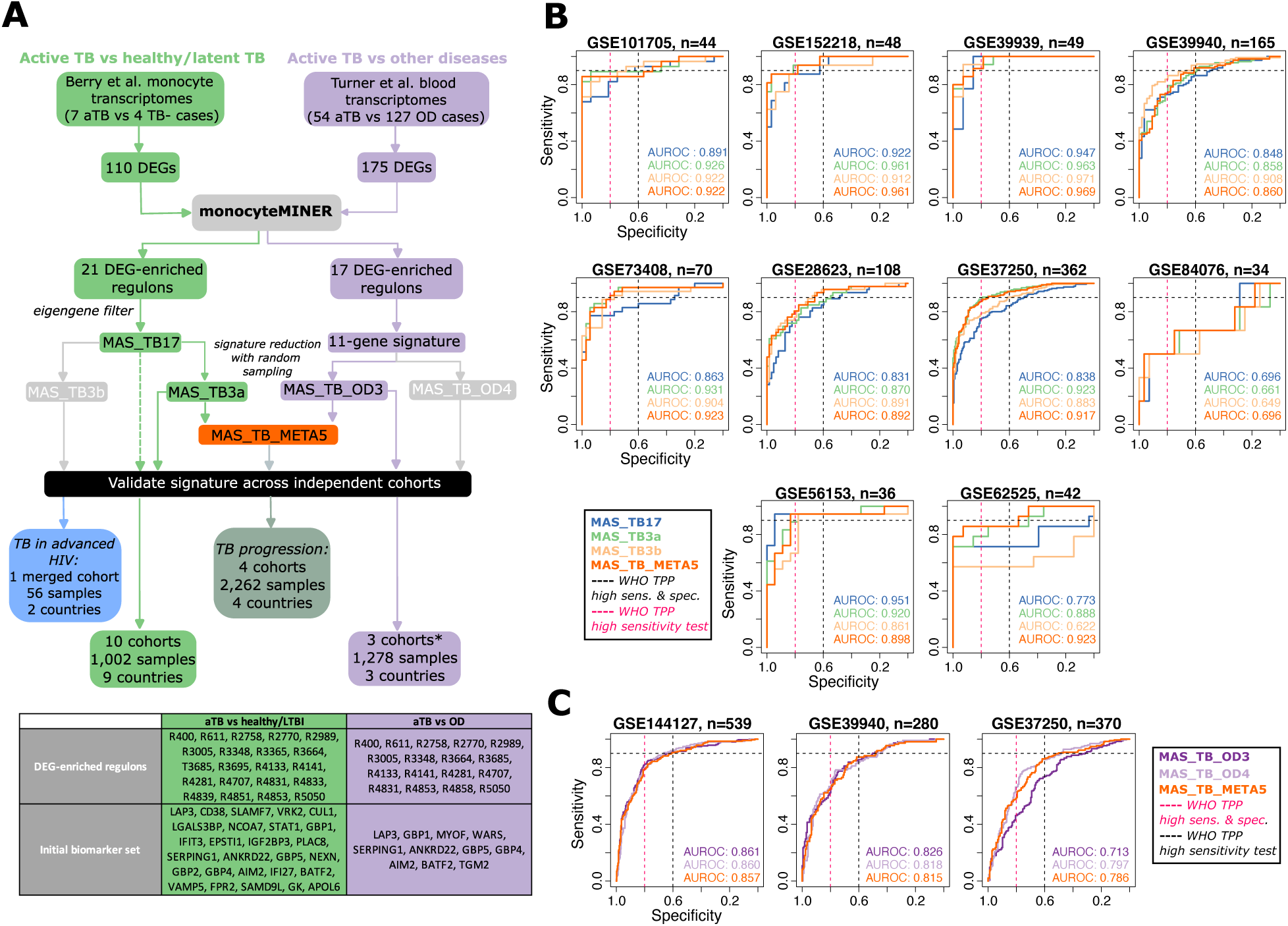
Discovery of monocyte-amplified transcriptional signatures for detection of active tuberculosis. A) Workflow used for discovering the MAS_TB17, MAS_TB3a and MAS_TB_OD3 signatures. The last two signatures were specifically defined for detecting aTB versus healthy controls (with or without a LTBI) and aTB vs. other diseases, respectively. General characteristics of cohorts used to retrospectively evaluate the performance of the proposed TB signatures across multiple disease contexts is also shown. The star (*) symbol indicates the use of two additional public datasets (from Turner et al.^36^ and Anderson et al.^45^) to evaluate performance of TB signatures not discovered using said cohorts for differentiating aTB from OD. B) Performance of MAS_TB17, MAS_TB3a, MAS_TB3b, and MAS_TB_META5 across 10 independent validation cohorts for aTB vs healthy/LTBI controls (Table 1). The number of samples analyzed from each cohort are indicated in the corresponding subpanel heading. Performance of each signature was measured using AUROC (values shown in each subpanel) using the PLAGE score implemented with TBSignatureProfiler. C) Performance of MAS_TB_OD3, MAS_TB_OD4, and MAS_TB_META5 across three independent validation cohorts for aTB vs. OD controls (Table 1). The number of samples analyzed from each cohort are indicated in the corresponding subpanel heading. Performance was evaluated as described for *panel B*.

**Table 1.**
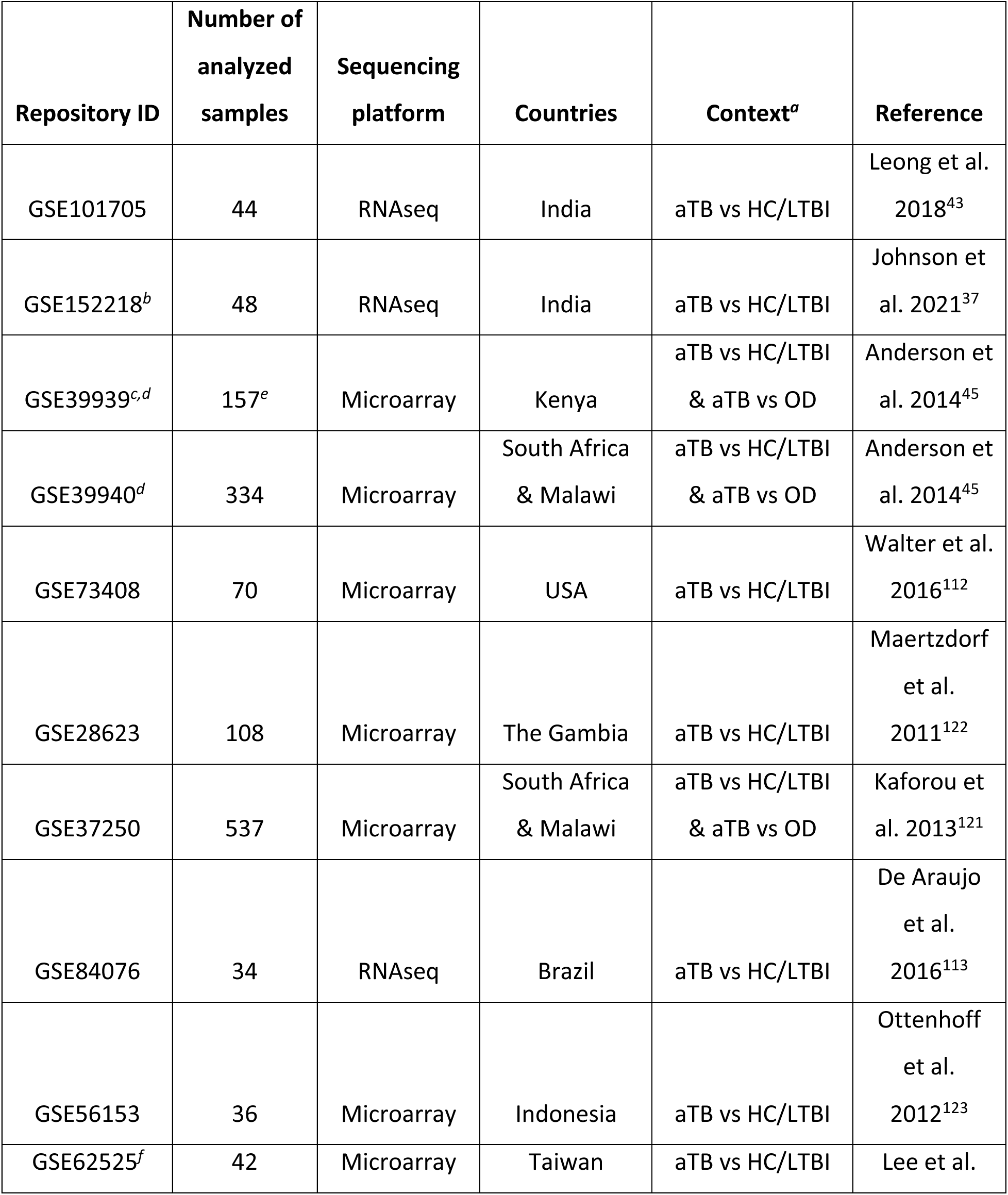
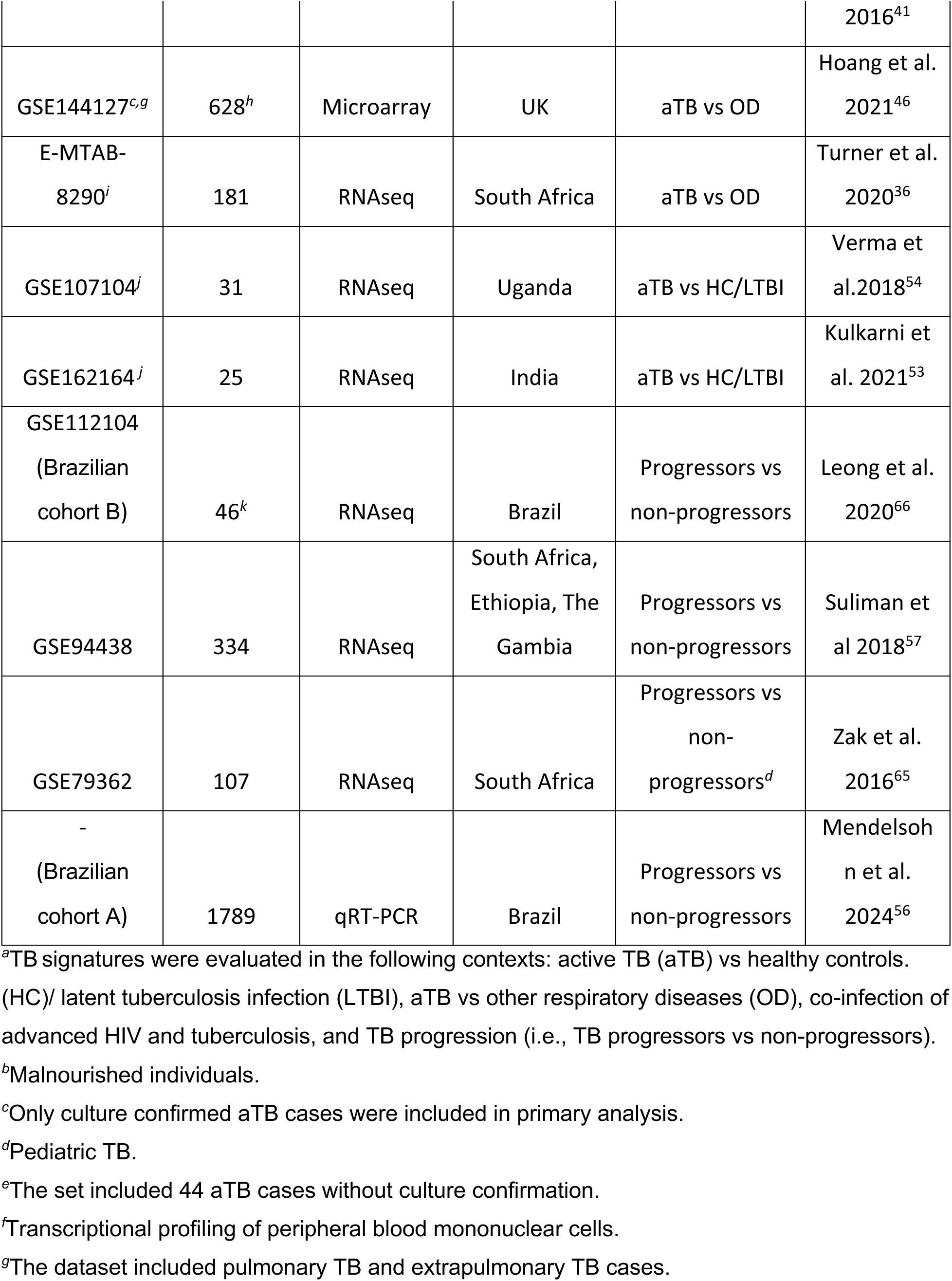
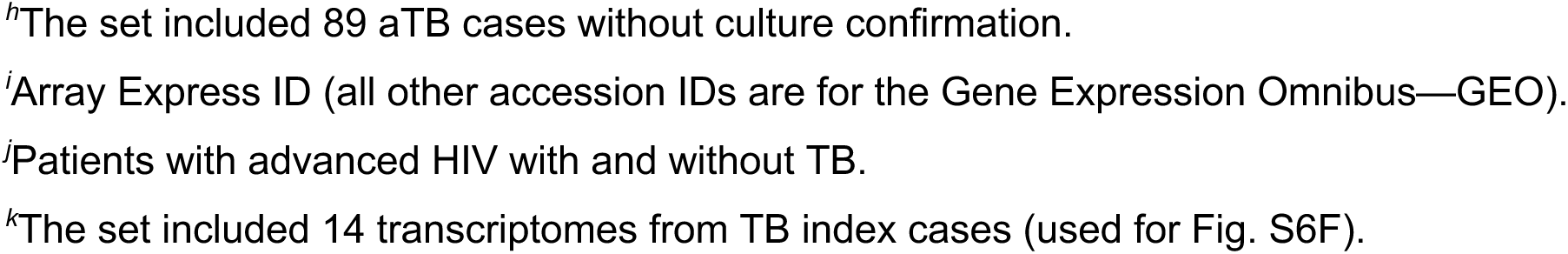
Cohorts used to compare and evaluate performance of TB signatures discovered in this study.

First, we discovered that 110 DEGs that differentiated monocyte transcriptomes of seven confirmed aTB cases relative to four healthy controls^35^ were enriched in 21 regulons within monocyteMINER. Through t-test comparisons of eigengenes for each of the 21 regulons, we identified 17 genes (here onward called MAS_TB17) from two regulons—R3664 and R3685—that were significantly differentially expressed across patients with aTB and controls (**Fig. 2A**; see ***Methods***). Performance of MAS_TB17 was evaluated on ten independent cohorts (i.e., cohorts that were not used for discovery) containing a total of 958 samples from nine countries across Africa, Asia and The Americas with 504 cases of aTB and 454 healthy controls/LTBI (**Table 1** and **Fig. 2B**). TBSignatureProfiler^37^ implementation of the pathway level analysis of gene expression (PLAGE) score^38,39^ demonstrated that MAS_TB17 had high sensitivity and specificity in detecting aTB across the independent cohorts with an average AUROC value of 0.856 (AUROC range: 0.696-0.951, **Fig. 2B**).

Benchmarking MAS_TB17 against four prior signatures discovered with comparably sized cohorts (Sambarey16^40^, Lee_4^41^, Verhagen_10^42^, Leong_24^43^) across 14 cohorts demonstrated superior performance in the aTB vs. OD context (75% of comparisons) and equivalent or better performance vs. healthy/LTBI controls (85% of comparisons), with particular advantage in advanced HIV patients (**Extended Data**, **Table S2**). Reducing MAS_TB17 to three-gene combinations by bootstrapped AUROC optimization (1,000 iterations, 50% random subsampling) identified two high-performing signatures: MAS_TB3a (ANKRD22, AIM2, VAMP5); and MAS_TB3b (IGF2BP3, AIM2, GBP5). Using a weighted AUROC^44^ (wAUROC, see ***Methods***) metric to account for cohort size, we determined that with wAUROCs of 0.900 and 0.877 for MAS_TB3a and MAS_TB3b, respectively, both signatures outperformed MAS_TB17 (wAUROC 0.849) in achieving WHO TPP across multiple independent cohorts (**Fig. 2B**, **Extended Data**, **Table S3**).

In parallel, we discovered 17 regulons within monocyteMINER that were enriched with overlapping subsets of 175 DEGs across 54 aTB patients relative to 127 patients with ODs (**Fig. 2A**; *see **Methods***). Sixteen out of the 17 regulons were also enriched with DEGs from the aTB vs healthy/LTBI comparison (**Fig. 2A**). Using methods described above, we identified an 11-gene TB signature (that included three genes, MYOF, TGM2 and WARS, that were unique to the aTB vs OD comparison, and eight genes that were shared with aTB vs healthy/LTBI analysis; **Fig. 2A**). The 11-gene signature was further refined and condensed with random sampling (n=1,000) using the original discovery cohort of 181 samples and an independent cohort of 35 aTB and 64 OD cases, respectively^36,45^ (**Table 2**). In so doing, we identified two four- (MAS_TB_OD4) and three- (MAS_TB_OD3) gene signatures (**Fig. 2A**) with statistically indistinguishable performance across two (of the three) independent validation cohorts (**Fig. 2C**). While both signatures achieved minimal TPP for a high-sensitivity screening test for aTB vs OD in one validation cohort, MAS_TB_OD3 achieved minimal TPP for a high-sensitivity screening test for aTB vs healthy/LTBI in 80% of the cohorts (compared to 50% for MAS_TB_OD4). Therefore, we selected the shorter of the two gene signatures (i.e., MAS_TB_OD3) and combined it with MAS_TB3a to generate the five-gene MAS_TB_META5 signature (ANKRD22, AIM2, VAMP5, GBP5, TGM2; **Table 2** and **Fig. 2A**) as a candidate multipurpose biomarker panel. Performance of MAS_TB_META5 in the validation cohorts previously described are shown in **Fig. 2B**, **Fig. 2C** and **Data Set S2**. In general, the performance of MAS_TB_META5 was indistinguishable from MAS_TB3a and MAS_TB_OD3 in their corresponding discovery contexts (**Table S3**). Moreover, MAS_TB_META5 had better performance (i.e., higher wAUROC) in detecting aTB within cohorts of both healthy/LTBI and OD than MAS_TB3a and MAS_TB_OD3 (**Data Set S2**). MAS_TB_META5 achieved WHO minimal TPP for a high-sensitivity screening test in two additional cohorts (i.e., in 7 out of 10) than MAS_TB3a for aTB vs healthy/LTBI, while achieving minimal TPP in the same number of cohorts for the other types of screening tests. While MAS_TB3a did not achieve TPP in any of the aTB vs OD cohorts, MAS_TB_OD3 and MAS_TB_META5 achieved the minimum TPP for a high-sensitivity screening test in the largest validation cohort of 539 samples (GSE144127^46^). Hence, the combination of MAS_TB3a and MAS_TB_OD3 into MAS_TB_META5 synergistically improved the overall performance in detecting aTB in cohorts containing either healthy/LTBI or OD cases.

**Table 2.** TB signatures identified in this work.

| Signature name | Genes | Discovery context | Discovery cohorts (number of samples) | DEG-enriched regulons | Associated TFs |
| --- | --- | --- | --- | --- | --- |
| MAS_TB17 | LAP3<br>CD38<br>SLAMF7<br>VRK2<br>CUL1<br>NCOA7<br>STAT1<br>EPSTI1<br>IGF2BP3<br>PLAC8<br>SERPING1<br>ANKRD22<br>GBP5<br>GBP4<br>AIM2<br>VAMP5<br>SAMD9L | Active TB vs healthy/latent TB infection | GSE19943<sup>a</sup><br>(11) | 3685, 3664 | MAFB |
| MAS_TB3a<sup>b</sup> | ANKRD22<br>AIM2<br>VAMP5 |  | GSE19943<br>GSE19439<br>GSE19442<br>GSE19444<br>(158) | All 21 DEG-enriched regulons<sup>e</sup> | - |
| MAS_TB3b<sup>c</sup> | IGF2BP3<br>AIM2<br>GBP5 |  |  | 5050 | STAT4 |
| MAS_TB_OD3<sup>c</sup> | AIM2<br>GBP5<br>TGM2 | Active TB vs other diseases | E-MTAB-8290<br>GSE39939<br>(280) | 5050 | STAT4 |
| MAS_TB_OD4<sup>b</sup> | WARS<br>ANKRD22<br>GBP5<br>TGM2 | Active TB vs<br>other<br>diseases | E-MTAB-8290<br>GSE39939<br>(280) | All 17 DEG-enriched<br>regulons<sup>e</sup> | - |
| MAS_TB_META5<sup>d</sup> | ANKRD22<br>AIM2<br>VAMP5<br>GBP5<br>TGM2 | Both | GSE19943<br>GSE19439<br>GSE19442<br>GSE19444<br>E-MTAB-8290<br>GSE39939<br>(438) | - | - |

<sup>a</sup>Only 11 monocyte transcriptomes in the dataset were analyzed.
<sup>b</sup>The signature with highest average AUROC in random sampling analysis with 1,000 iterations (see methods).
<sup>c</sup>Selected individual regulon-based gene signatures (see methods).
<sup>d</sup>Signature for differentiating active TB from both healthy/latent TB infection cases and other diseases.
<sup>e</sup>Relevant genes are distributed across all DEG-enriched regulons identified for the TB context of interest (shown in Fig. 5A).

### Comparative performance assessment of MAS_TB_META5

We postulated that a multi-purpose TB signature should have minimal performance tradeoff in diagnosing aTB in two different contexts, i.e., differentiation from healthy/LTBI cases and differentiation from ODs. We compared the performance of the MAS_TB signatures (**Table 2)** and 34 TB signatures in TBSignatureProfiler that were comprised of ≤ 10 genes each, and, therefore, deemed most amenable for implementation into a diagnostic test in a clinical setting^5,44^ (**Extended Data**). Using the PLAGE score, we identified top performing TB signatures (ROC curve comparison p-value > 0.05 with respect to the signature with highest AUROC; **Fig. 3A)** in each of the ten independent cohorts for aTB vs healthy/LTBI cases, and five cohorts for aTB vs OD cases (after excluding for each TB signature any dataset used for its discovery or training; **Data Set S2**). For example, only three of the five cohorts comprising aTB vs OD cases, not used for discovery, were used as independent datasets for testing the performance of the two MAS_TB_OD signatures and MAS_TB_META5 (see ***Methods***).

**Figure 3.**
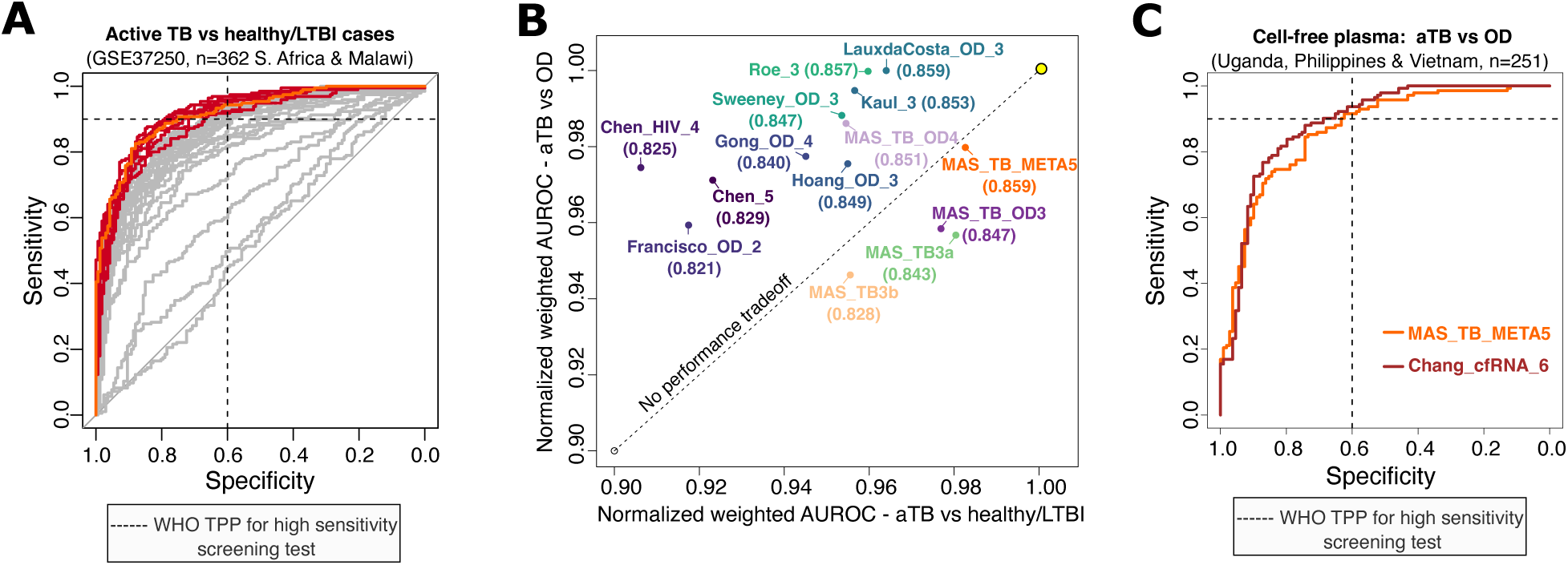
Comparative assessment of MAS_TB_META5, as a multipurpose TB signature. A) Performance of all concise TB signatures (i.e., with 10 or less genes) evaluated in the validation cohort by Kaforou et al.^121^ for aTB vs LTBI. The GEO series ID, total number of samples included in the analysis, and the countries where samples were collected are shown in parenthesis. As explained in the main text, AUROCs for the ten top performing signatures (dark shades of red), were statistically indistinguishable from each other. MAS_TB_META5 ROC curve is shown in orange color. B) MAS_TB_META5, comprised of the union of MAS_TB3a and MAS_TB_OD3 gene sets (Fig. 2A), outperformed previous TB signatures at differentiating active TB from both healthy controls and other diseases. Overall performance of each signature in differentiating aTB from healthy controls and LTBIs (x-axis) and other diseases (y-axis) across all respective validation cohorts was integrated into a single weighted mean AUROC score (wAUROC), as described in the main text. The scores were normalized with respect to the maximum wAUROC among all evaluated TB signatures. Only top performing signatures in at least two validation cohorts for aTB vs OD (in addition to MAS_TB3a) are shown. The number in parenthesis for a given signature indicates its wAUROC score computed across all validation cohorts (x- and y-axis), excluding datasets used for its discovery or training. Signatures with no performance tradeoffs across the two comparisons would be expected to fall on the dotted diagonal line with the ideal performance indicated by the yellow dot in the top right corner. C) Performance of MAS_TB_META5 in differentiating aTB from OD in cell-free plasma-derived transcriptomes^47^. As a reference, the performance of the 6-gene signature originally discovered using the plasma transcriptomes (i.e., Chang_cfRNA_6) is also shown as the red curve.

To objectively evaluate the multipurpose utility of all TB signatures, we used the wAUROC metric to quantify performance tradeoffs in detecting aTB across the two contexts, i.e., vs. healthy/LTBI and vs. ODs^44^. With the highest wAUROC (0.859), MAS_TB_META5 outperformed all 34 TB signatures at differentiating aTB from both healthy/LTBI and ODs, with no performance tradeoffs (**Fig. 3B**), across all independent validation cohorts, even when suspected TB cases that were not culture-confirmed in the GSE39939 and GSE144127 datasets were included in the analysis **(Fig. S2**; *see **Methods***). Notably, MAS_TB_META5 achieved WHO TPP for a high sensitivity screening test for detecting extrapulmonary TB in the GSE144127 dataset (**Fig. S3**), as well as for a high-sensitivity screening test for differentiating aTB from ODs in a multi-country compendium of transcriptomes from cell-free blood plasma^47^ (**Fig. 3C**). While a 6-gene signature, originally discovered using the same cell-free plasma dataset, had comparable performance as MAS_TB_META5, it was unable to distinguish confirmed aTB cases from OD in the UK cohort (GSE144127)^46^ (**Fig. S4**). In summary, these findings demonstrate that MAS_TB_META5 was the best performing signature with the least performance tradeoff in detecting aTB among healthy, LTBI and OD cases across 13 independent cohorts of ∼2,150 patients from 10 countries and 5 continents.

### MAS_TB_META5 meets WHO TPP for detecting active TB in patients with advanced HIV

ART initiation can cause a lethal TB-associated immune reconstitution inflammatory syndrome (TB-IRIS) in TB co-infected HIV patients^48^, due to hyperactive response of the recovering immune system to TB antigens, causing or worsening TB symptoms^49,50^. TB testing is therefore important before initiation of ART, especially in advanced HIV patients, who are at significantly higher risk of developing TB, due to a compromised immune system with < 200 CD4^+^ T cells per μL of blood^51,52^. However, transcriptional signatures discovered through training on aTB cases vs. healthy/LTBI/ODs are suspected to incur performance tradeoff in detecting TB in ART-naive HIV-co-infected patients, due to higher baseline expression of interferon-related genes in these patients^50^. Co-infections with other diseases may also reduce performance of TB signatures due to cross-reactivity^50^. While Kulkarni2^53^ and Verma2^54^, discovered using cohorts of advanced HIV patients from Uganda and India with and without TB co-infection (**Table 1**), expectedly performed significantly higher in these datasets, their performance is yet to be evaluated on independent cohorts of advanced HIV patients with TB.

Given these challenges and the urgent need for a test to detect aTB in advanced HIV patients, we evaluated the performance of MAS_TB signatures (**Table 2**) and the 34 prior signatures in detecting 29 confirmed aTB cases among 56 advanced HIV patients in a combined India/Uganda cohort (see ***Methods*** for details on how data from the two cohorts were combined). The Kulkarni2 signature was excluded from this analysis as it was discovered using one of these datasets. All five MAS_TB signatures were among the top performers with statistically indistinguishable AUROCs (e.g., MAS_TB3a AUROC was 0.922 with 95% CI: 0.849-0.995; and MAS_TB_META5 AUROC was 0.911; with 95% CI: 0.824-0.998; **Fig 4A**). The PLAGE score for MAS_TB_META5 in TB-HIV co-infected patients was significantly different from that of HIV patients who did not have aTB (**Fig. 4B**; t-test p-value = 8.5e-10). The same trend was observed for the previously reported TB signatures (**Fig. S5**). Most importantly, MAS_TB META5, MAS_TB3a, MAS_TB3b, and MAS_TB_OD3 achieved WHO TPP for a high-sensitivity TB screening test in advanced HIV patients (**Fig. 4C**). Among the eight (out of the 34 previous TB signatures) that also achieved WHO TPP, five had potential multipurpose utility (shown with bold font in **Fig. 4A**, **Data Set S2)**. Importantly, MAS_TB_META5 approached the WHO TPP for a high-sensitivity and high-specificity screening test with specificity of 88.9% at a sensitivity of 89.6%. Notwithstanding the relatively small size of the combined dataset, our findings demonstrate that the aTB signal detected by MAS_TB_META5, as well as MAS_TB3a and MAS_TB3b, is not dampened in advanced HIV patients.

**Figure 4.**
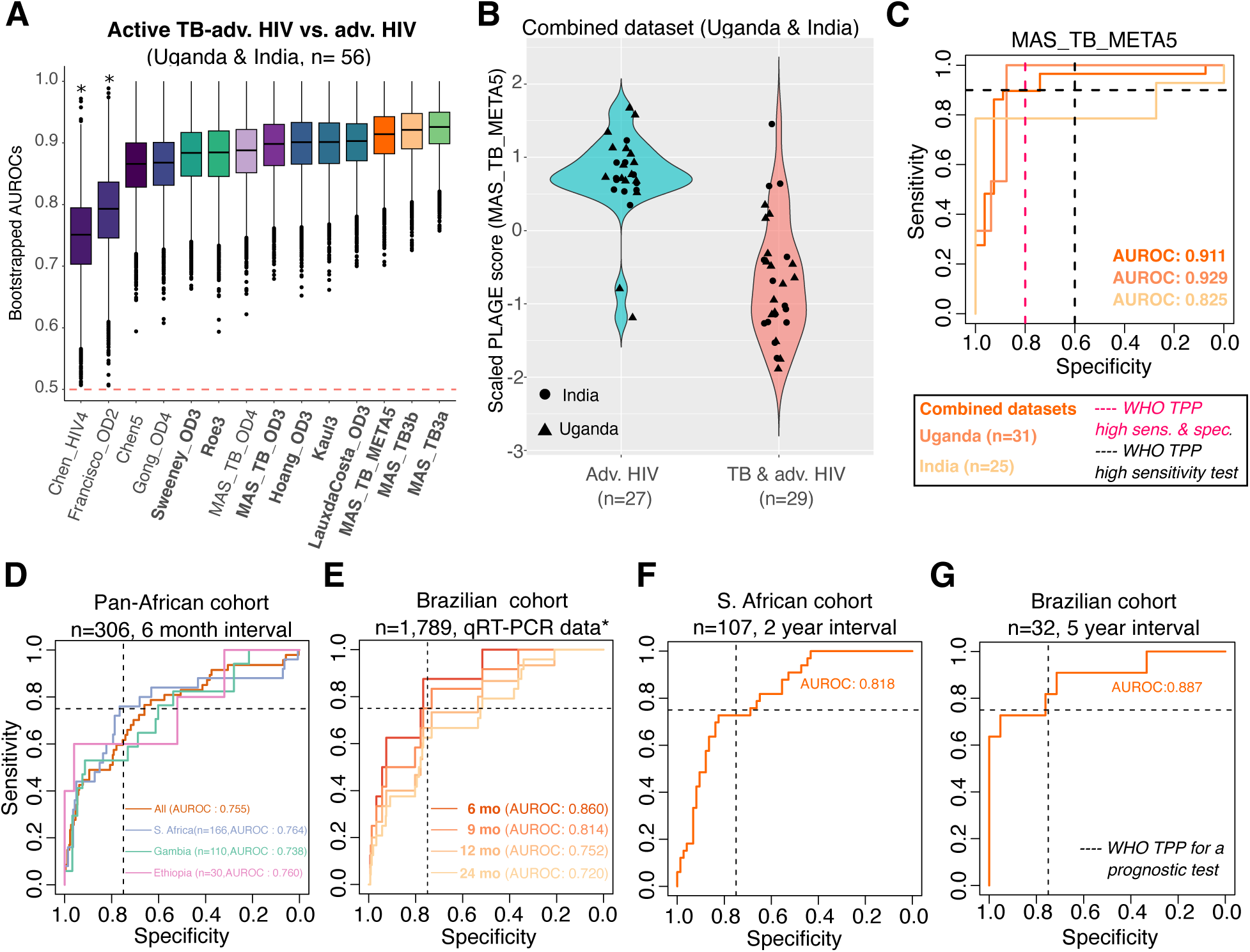
The MAS_TB_META5 signature achieves WHO minimum recommended performance for a high-sensitivity TB screening test in advanced HIV populations and for a prognostic test in patients in contact with active TB cases. A) Comparison of performance of gene signatures shown in Fig. 3B in the combined advanced HIV cohort. Two small cohorts, from Uganda (GSE107104) and India (GSE162164; Table 2), were combined into a single cohort as explained in the Methods. Boxplots show the distribution of AUROC for each signature across 10,000 bootstraps (see Methods). Signatures are ranked by median AUROC. Signatures that achieved WHO TPP for a high-sensitivity screening test (sensitivity ≥ 90% and specificity ≥ 60%) are shown in bold font. The red horizontal dashed marks the expected AUROC of a random classifier. B) Violin plot with the z-score transformed PLAGE scores for the MAS_TB_META5 signature for advanced HIV patients with and without TB co-infection before start of antiretroviral treatment. C) ROC curves for MAS_TB_META5 in the combined cohort described above, and in the individual cohorts. D) Performance of MAS_TB_META5 at predicting patients that would progress to active TB in a cohort of patients from South Africa, The Gambia and Ethiopia (GSE94438). ROC curves for a cohort with 47 TB progressor samples and 259 non-progressor samples from three countries (and per country) are shown for a six-month interval defined as the time elapsed between pre-disease sample collection and TB diagnosis (as explained in the Methods). The first collected sample of each non-progressor was used as control. E) Prognostic performance of MAS_TB_META5 in predicting progression to aTB using qRT-PCR profiles of target transcripts across 1,789 patients in a Brazilian cohort^56^ (‘Brazilian Cohort A’). The “*” symbol indicates that the mRNA levels of only three genes (ANKRD22, GBP5 and VAMP5) out of the five in MAS_TB_META5 were measured in this study. To facilitate comparison with signatures previously evaluated with this dataset, time intervals were defined as the time elapsed between enrollment (i.e., first sample collection) and TB diagnosis. To evaluate the potential of clinical implementation of MAS_TB_META5, the signature score was defined as the sum of the delta Ct values of the three genes present in the dataset. F) Performance of MAS_TB_META5 in the Adolescent Cohort Study^65^ (ACS). For each individual among the 33 TB progressors and 77 non-progressors, signature scores were computed using their initial sample. G) Performance of MAS_TB_META5 in a cohort of 32 patients from Brazil (‘Brazilian Cohort B’) followed for five years (11 patients progressed to active TB during the study; GSE112104). Unless mentioned otherwise, all results correspond to PLAGE scores.

### MAS_TB_META5 predicts risk of progression to active TB across multiple cohorts

A prognostic test to predict likelihood of progression to aTB in high-risk individuals, living in households with aTB patients or regularly exposed in a TB endemic region, can potentially curtail transmission of the disease by enabling early detection and treatment^55^. Although WHO has recommended a two-year interval for predicting TB progression (i.e., time elapsed between prediction and aTB diagnosis), there is evidence that host transcriptional biomarkers may be better suited for predicting progression at shorter intervals^5,56^ while still offering benefits for TB treatment programs^56^. Hence, we evaluated the prognostic performance of MAS_TB_META5 across four cohorts of TB progressors and non-progressors from Brazil, South Africa, The Gambia and Ethiopia, which included confirmed progression events at 6 (pan-African cohort and Brazilian cohort A), 9 (Brazilian cohort A), 12 (pan-African cohort and Brazilian cohort A), 24 (pan-African cohort, adolescent S. African cohort and Brazilian cohort A), and 60 months (Brazilian cohort B) (**Table 1**).

None of the signatures in TBSignatureProfiler, including Sulliman4^57^, which was discovered using the same cohort (i.e., GSE94438^57^), achieved the WHO TPP for a prognostic test (i.e., specificity ≥ 75% at sensitivity ≥ 75%^58^) at the 6-month or longer intervals for the combined African cohort of 306 participants from S. Africa, The Gambia, and Ethiopia^57^ when evaluated using the PLAGE score. Although the underlying reasons are yet to be determined, consistent with findings from previous African multi-country studies^59^, we observed variability in the performance of TB signatures across countries. While MAS_TB_META5 did not achieve WHO benchmarks in cohorts from The Gambia (n=110) and Ethiopia (n=30), which also reduced its performance in the full African cohort, it did meet the WHO minimal TPP for a six-month interval for the S. African cohort of 166 participants (AUROC: 0.764; 95% CI: 0.641-0.886; p-value= 0.0016) (**Fig. 4D**). In addition to MAS_TB_META5, WHO TPP for prognosis was achieved for the S. African cohort by seven previously reported signatures, of which only Gliddon_OD3^60^, Jacobsen3^61^ and Gong_OD_4^62^ had < 10 genes. Notably, the performance of MAS_TB_META5 at the two-year interval also approached WHO minimal TPP for the S. African cohort (AUROC: 0.710; 95% C.I: 0.611-0.809; p-value: 0.0009; **Fig. S6A)** with a sensitivity of 69.2% at a specificity of 70.2%. MAS_TB_META5 also approached WHO TPP in the 6- and 12-month interval in the full cohort (**Fig. S6B**).

Mendelsohn et al. monitored a cohort of 1,789 healthy patients (“Brazilian cohort A”, **Table 1**) for aTB at 6-,9-, 12-, and 24-month intervals after their whole blood was transcriptionally profiled^56^. However, this study used qRT-PCR to quantify transcripts encoded by a set of 80 genes, which included just three of the five genes (ANKRD22 and VAMP5 from MAS_TB3a, and GBP5 from MAS_TB_OD3) in the MAS_TB_META5 signature. Notwithstanding this limitation, the predicted risk score for progression to aTB calculated using ΔCt values (see ***Methods*** and Mendelsohn et al.^56^) for the three genes in MAS_TB_META5 achieved WHO TPP up until the 9-month interval (from time of enrollment) with a maximum specificity of 77.9% at sensitivity ≥ 75% (6-month AUROC: 0.86, 95% C.I: 0.745-0.974, p-value: 0.0081; 9-month AUROC: 0.814, 95% CI: 0.699-0.929, p-value: 0.0039; **Fig 4E**). The performance of this prognostic assessment was unchanged when the risk for progression to aTB was estimated using the sum of Ct values (**Fig. 4E**) or with the PLAGE score (**Fig. S6C**). Notably, only three previous TB signatures (Gliddon4^1^, Rajan5^63^ and Duffy9^64^), but none that made the cut for multipurpose utility (**Fig. 3B**), achieved WHO TPP at the 9-month interval^56^. MAS_TB_META5 outperformed all three of these signatures, as the only signature with an AUROC ≥ 0.85, regardless of how the 6-month interval was defined, either from enrollment time (**Fig. S6D**) or the last time point when the transcriptome was profiled before disease diagnosis (as done for the pan-African cohort; **Fig. S6E**).

Next, we evaluated the performance of MAS_TB_META5 in a S. African cohort of 104 adolescents (ages 12-18 years; 74 non-progressors and 33 TB progressors), who were followed up to 24 months for progression to aTB (**Fig. 4F**). Of the 33 TB progressors in this study, the baseline blood transcriptomes of 23 patients were profiled on day 0, eight were profiled on day 180, and two at day 360^65^ (*see **Methods***). Similarly, among the 74 non-progressors, two were initially profiled on day 180 and two on day 360^65^. The performance of MAS_TB_META5 was lower (AUROC: 0.789, 95% CI: 0.691-0.886, p-value: 0.0007) when only 23 progressors and 70 non-progressors with day zero 0 transcriptome profiles were considered. However, when the entire set of blood transcriptomes of non-progressors and progressors were considered, the performance of MAS_TB_META5 for the two-year interval approached WHO TPP guidelines with maximum sensitivity of 72.7% at specificity ≥ 75% (AUROC: 0.818, 95% C.I: 0.736-0.899, p-value = 3.41e-06).

Finally, we evaluated MAS_TB_META5 performance in predicting risk of TB progression at five-year interval in a second Brazilian cohort of 32 patients (11 progressors and 21 non-progressors; “Brazilian cohort B”, **Table 1**)^66^. PLAGE scores for MAS_TB_META5 across the non-progressors was significantly higher relative to index cases (T-test p-value = 0.0015), whereas scores for TB progressors were indistinguishable from the index cases (**Fig. S6F**). Notably, MAS_TB_META5 achieved WHO TPP for a prognostic test by predicting TB progression at a five-year interval with a maximum sensitivity of 81.8% at specificity ≥ 75% (**Fig. 4G**). Together these findings demonstrate through evaluations on cohorts from different countries that MAS_TB_META5 consistently achieved or approached WHO TPP for a prognostic test to predict risk of progression to aTB at 6-months up to a 5-year interval.

### Monocyte-amplified gene signatures are mechanistically associated with diverse aspects of human-pathogen interactions in active pulmonary TB

Next, we examined the underlying reason for the consistently high performance of the MAS_TB signatures across contexts by investigating their association with biological processes relevant to TB infection. The MAS_TB signatures were derived from an active TB network comprising 110 genes (including the 30 DEGs in **Fig. 5A**) organized across 22 regulons—13 enriched with monocyte-specific genes and 16 enriched with interferon-related genes^67^ (*see **Methods***)—regulated by 16 TFs that revealed a coherent picture of human-Mtb interaction during aTB. The biological relevance of this network is evident in the extensive prior associations of its components with TB pathogenesis. At least 11 of the 16 TFs in this network have established roles in TB biology (**Fig. 5A** and **Data Set S3**). For instance, single nucleotide polymorphisms in MAFB, which regulates regulons R3664 and R3685, the source for MAS_TB17, have been linked to increased TB risk in human populations^68,69^. MAFB’s dual role in monocyte-to-macrophage differentiation^70^ and macrophage response to Mtb^71^ underscores its mechanistic importance. Similarly, STAT4, the transcriptional regulator of R5050 from which MAS_TB3b emerged, is essential for Mtb clearance in animal models^72^. This functional and mechanistic association with TB infection and clearance was observed across 25 out of 30 genes in the aTB network (**Data Set S3**).

**Figure 5.**
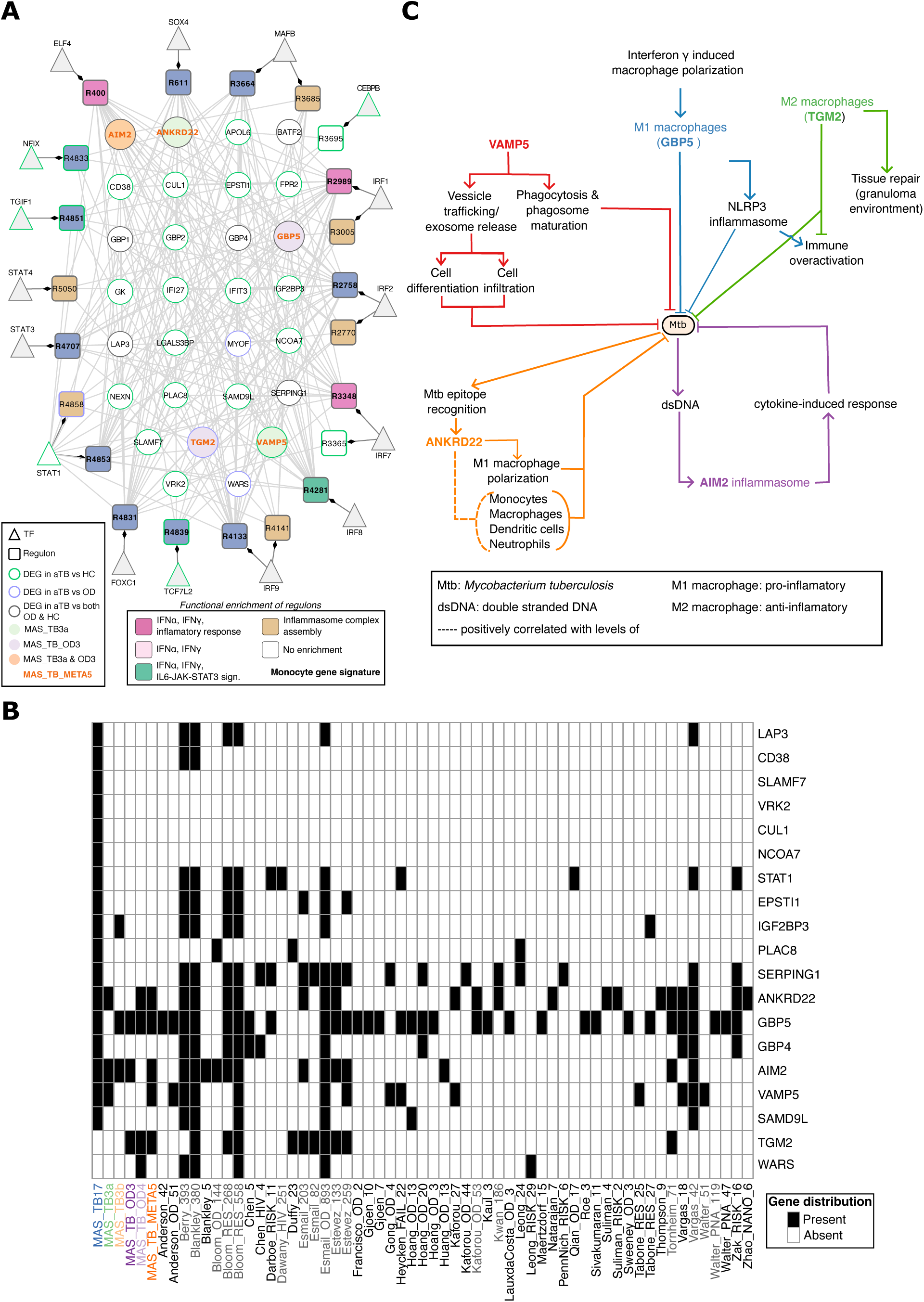
The TB-dysregulated transcriptional network offers insights into the host defense strategies against *M. tuberculosis*. A) A monocyteMINER subnetwork of 16 TFs regulating 110 genes across 22 regulons enriched for DEGs (hypergeometric test p-value < 0.05) in aTB patients relative to healthy/LTBI controls or ODs (Fig. 2A). To facilitate visualization, only 30 DEGs within these regulons are displayed. Nodes depicting unique genes that comprise MAS_TB3a and MAS_TB_OD3 are shown in green and purple, respectively, and AIM2, a gene common to MAS_TB3a and MAS_TB_OD3, is shown as an orange node. Regulons were color-coded based on functional enrichment of MSigDB hallmarks; GO terms are shown where there was no enrichment of hallmarks. Regulons enriched for monocyte markers and genes in MAS_TB_META5 are shown in bold font. Refer to figure key for interpreting color-code for node borders. B) Composition of genes across the five MAS_TB signatures reported in this study and 55 previously reported TB signatures in TBSignatureProfiler that share genes within at least one MAS_TB signature. The nomenclature for gene signatures, borrowed from TBSignatureProfiler, is a combination of the last name of the first author of the manuscript reporting the signature followed by an underscore and the number of genes reported in said signature. TB signatures with >41 genes (i.e., the top 25% of all signatures in TBSignatureProfiler) are shown in grey font. C) Genes in MAS_TB_META5 are associated with five distinct host immune defense mechanisms triggered during active TB.

While MAS_TB17 shares some genes with the 77 TB signatures in TBSignatureProfiler^37^, four genes (SLAMF7, VRK2, CUL1, and NCOA7) are unique and absent from all prior signatures. More importantly, the specific gene combinations of all MAS_TB signatures are distinct from previously reported signatures of comparable length (**Fig. 5B**). Statistical analysis confirms the novelty of these combinations. The probability of randomly selecting genes matching MAS_TB3a and MAS_TB3b compositions, where constituent genes appear in at least 13 and 6 of the 77 existing signatures, respectively, was exceptionally low (hypergeometric test p-values: 1.7e-07 and 0.00011). This indicates that the two gene MAS signatures could not have been derived simply from analyzing existing signature compositions. For context, MAS_TB3b contains genes with varying prevalence across the 77 signatures: GBP5 (present in 32 signatures), AIM2 (13 signatures), and IGF2BP3 (6 signatures). Given that 115 genes appear in 6 or more existing signatures, MAS_TB3b represents just one of 246,905 possible three-gene combinations. Similarly, while MAS_TB_OD3 and MAS_TB_OD4 include genes (e.g., GBP5 and ANKRD22) also present across multiple existing TB signatures, they also include genes (e.g., TGM2 and WARS) that were not part of any prior signature composed of < 23 genes (**Fig. 5B**). Similarly, MAS_TB_META5 genes are present in at least 10 of the 77 previous TB signatures considered in this analysis. Thus, MAS_TB_META5 is one of 324,632 possible 5-gene combinations among the 35 genes present in at least 10 of the previous TB signatures.

The biological comprehensiveness of MAS_TB_META5 illustrates why monocyteMINER-derived signatures may capture disease states with high sensitivity and specificity across diverse contexts. The five genes in MAS_TB_META5 (AIM2, GBP5, ANKRD22, TGM2, and VAMP5) capture five layers of host immune defense against Mtb, playing distinct but interrelated roles spanning vesicular trafficking, inflammasome activation, granuloma maintenance, myeloid cell polarization, and cytosolic DNA sensing (**Fig. 5C**). **VAMP5**, a SNARE-family vesicle trafficking protein^73,74^, facilitates phagosome formation, an early and necessary step that initiates the downstream maturation cascade through which phagolysosomes develop their acidic, microbicidal environment^74,75^. Notably, Mtb is known to evade this mechanism of killing precisely by blocking phagosome maturation and its characteristic acidification^75^. **GBP5** (Guanylate Binding Protein 5) is an interferon-gamma (IFN-γ)-inducible gene that serves as a critical activator of NLRP3 inflammasome assembly in pro-inflammatory M1 macrophages^76,77^. Because GBP5 sits at the intersection of IFN-γ signaling and NLRP3-mediated innate immune activation^77^, both of which are hallmarks of the macrophage-driven inflammatory state during active Mtb infection^78,79^, its transcription in whole blood is sharply and consistently upregulated in aTB relative to LTBI and healthy controls across multiple independent cohorts^80^. **TGM2** (Transglutaminase 2) is a conserved marker of alternatively activated, anti-inflammatory M2 macrophages in both humans and mice^81^. Although both M1 and M2 macrophages reside in the lungs of TB patients, in granulomatous zones the M2-polarized macrophages are most prevalent, and are associated with dampened bactericidal activity and conditions permissive for Mtb persistence^82^. This explains why Mtb actively induces M2-polarization through virulence factors such as ESAT-6^83^. TGM2 activity promotes Mtb intracellular survival in macrophages through autophagy-dependent mechanisms, and its pharmacological inhibition limits bacterial replication *in vitro* and decreases bacterial burden *in vivo*^84,85^. **ANKRD22** (Ankyrin Repeat Domain-containing Protein 22) is an N-myristoylated protein^86^ whose expression is induced in activated monocytes and M1 macrophages^87^. In whole-blood transcriptome data from active TB patients, ANKRD22 expression is positively correlated with the abundance of myeloid immune cell types (e.g., monocytes, macrophages and neutrophils) with expanded populations in active relative to latent TB^88^. Mechanistically, ANKRD22 induces M1 macrophage polarization by increasing phosphorylation levels of the transcriptional factor Interferon Regulatory Factor 3 (IRF3) through its interaction with the mitochondrial antiviral-signaling protein (MAVS), thereby driving the transcription of IFN-β and related genes^87^. Finally, **AIM2** (Absent In Melanoma 2) senses double-stranded DNA^89^ from bacterial pathogens, including Mtb, within the macrophage cytosol to drive the inflammasome assembly, triggering activation of caspase-1 and secretion of interleukin 1β (IL-1β) and interleukin 18^90^; however, its contribution to IL-1β release in human macrophages remains contested^91^. AIM2-deficient mice show impaired defense against Mtb infection and defective Th1 responses (e.g., IFN-γ)^90,91^. Notably, virulent Mtb leverages its ESX-1 secretion system to suppress AIM2 activation, which likely represents a specific immune subversion strategy in the host-pathogen arms race^92^. These findings show that monocyteMINER identified MAS signatures from functionally interconnected gene modules, avoiding redundant selection of multiple genes from the same functional module. In so doing, the modular TRN-based approach enabled comprehensive selection of unique genes that collectively encode the full spectrum of host-pathogen interactions, offering a plausible explanation for the robust performance of MAS_TB signatures across diverse populations and disease contexts.

## DISCUSSION

This study introduces a novel systems biology approach to biomarker discovery by leveraging cell type-specific mechanistic regulatory networks to decode disease signals from whole blood transcriptomes. Rather than treating blood as an undifferentiated mixture of signals, the monocyte-centric approach developed in this study recognizes that these circulating sentinel cells serve as an immunological surveillance system, continuously sampling tissue environments and internalizing information about systemic health status. Constructed using monocyte transcriptomes from 1,202 ethnically-diverse healthy individuals, monocyteMINER is a comprehensive TRN model of 10,567 genes organized into 5,682 functionally coherent regulons, that leverages this extravascular surveillance function of monocytes to serve as a biological lens through which disease-perturbed signals can be amplified and mechanistically interpreted.

The monocyteMINER framework addresses fundamental limitations of conventional biomarker discovery approaches that have yielded disappointingly few clinically implemented tests despite decades of effort^93,94^. The power of this approach stems from its biological grounding, and, by extension, the mechanistic association of the biomarkers it identifies with disease etiology and progression^12^. Because of the ‘curse of dimensionality’ of multiomics datasets that has plagued biomarker discovery^95^, traditional ML methods typically identify statistical correlations without explicitly taking into account the underlying biology^12^. In contrast, using principles of systems biology, monocyteMINER organizes genes into functionally coherent regulons based on evidence of mechanistic co-regulation by specific transcription factors. This modular organization ensures that selected biomarkers represent distinct biological processes rather than redundant measurements of the same pathway. For TB, this meant capturing the full spectrum of host-pathogen interactions, from bacterial DNA recognition and inflammasome activation to macrophage polarization and intercellular communication, within just five genes (MAS_TB_META5; **Fig. 5C**). The biological coherence of MAS_TB_META5 explains its consistent high-level performance across 12 countries, multiple RNA profiling platforms, sample types (whole blood, peripheral blood mononuclear cells, and plasma transcriptomes), and diverse clinical contexts, including advanced HIV co-infection and prediction of TB progression.

The efficiency of biomarker discovery achieved through monocyteMINER is particularly striking. While conventional approaches required cohorts of thousands of patients to identify robust TB signatures, exemplified by Sweeney et al.’s use of 1,023 samples across 3 datasets to derive their 3-gene signature^80^, the monocyteMINER approach used a modest size training set of 438 subjects (across four small cohorts) to discover a five-gene MAS_TB_META5 signature that outperformed 77 existing signatures (**Fig. 3B** and **Fig. 6**). This efficiency stems from the network-enabled amplification of disease-relevant signals, that builds on the transcriptional programs most perturbed in monocytes during disease. Although network-based strategies, mainly leveraging correlation, pathway analysis and protein-protein interaction networks, have been fruitful in the past for biomarker discovery^21,35,40,62,96^, our study went beyond previous efforts by reconstructing a cell type-specific mechanistic hypothesis-generating model. While these findings align with emerging evidence that cell type-specific approaches outperform bulk transcriptomics for biomarker discovery^97,98^, what we have shown is that the cell-type specific lens is just needed for discovery of highly sensitive and specific transcriptional biomarkers from monocyte or whole blood transcriptomes, and, more importantly, that once discovered the signature can be subsequently detected in whole blood or plasma. The atherosclerosis findings further illustrate this principle: despite using an imperfect proxy (CAC scores) for disease risk, the atherosclerosis-perturbed network revealed mechanistic insights into inflammatory processes driving atherosclerosis that translated into diagnostic markers for both CAD and AMI.

**Figure 6.**
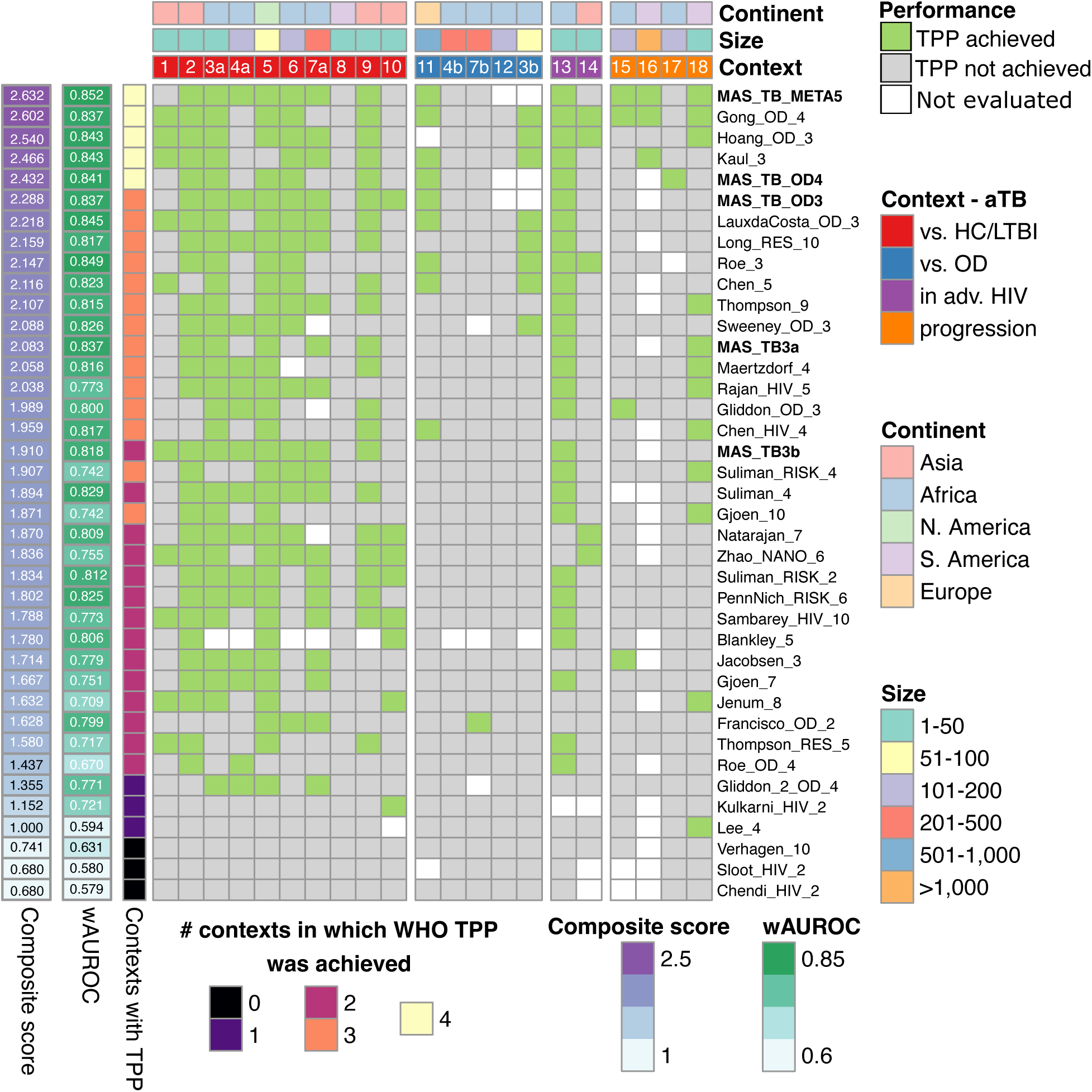
MAS_TB_META5 outperforms other TB signatures in achieving WHO TPP for a high-sensitivity screening and prognostic test across contexts. Eighteen cohorts used for validation of signature performance are organized into four contexts (see “<u>Context</u>” track), with numbers of subjects in each cohort indicated in “<u>Size</u>” track, and geographic location noted in the “<u>Continent</u>” track. For datasets that contain aTB cases, OD cases, and healthy/latent TB infection cases, “a” indicates performance assessment in aTB vs. healthy/LTBI context, and “b” indicates assessment in aTB vs.OD context. Signatures are ranked based on a “<u>Composite score</u>”, calculated as the sum of normalized values of “<u>wAUROC</u>” (to account for cohort size) and the number of cohorts and contexts in which WHO TPP was achieved (“<u>Context with TPP</u>”). MAS TBsignatures are shown in bold font. In the heatmap, green shading indicates cohorts in which WHO TPP was achieved, gray shading indicates cohorts where WHO TPP was not achieved, and white indicates cohorts used for discovery of the relevant signature, and hence excluded from its performance assessment. **<u>Notes</u>**: Dataset # 15 (GSE94438) was used to evaluate performance of each TB signature in a 6-month window for samples from South Africa, as explained in the main text and shown in Fig. 4D. Dataset # 16, which includes qRT-PCR profiles of 80 genes, was used to evaluate the performance of 20 TB signatures over a 9-month window from enrollment^56^. While the evaluation was performed using normalized scores published as part of the study, the star symbol (“*”) indicates evaluations performed by scoring the sum of delta Cts of partial or complete set of member genes within each signature (i.e., MAS_TB_META5, Hoang_OD_3 and Kaul_3 with three out of five, two out of three, and all three genes in the signatures, respectively). As recommended by the authors^62^, Gong_OD_4 was evaluated using the geometric mean of delta Ct values of three out of the four genes in the signature for which qRT-PCR profiles were available in this study. Dataset # 17 (GSE79362) was used to evaluate performance of each TB signature over a 24-month window. Dataset # 18 (GSE112104) was used to evaluate performance of each TB signature over a 5-year window.

The clinical implications of this approach extend far beyond the diseases studied here. Monocytes encounter and respond to diverse pathological processes, spanning infections and autoimmune disorders to cancers and neurodegenerative diseases^13–17,22^, making them ideal cellular reporters of systemic health. The monocyteMINER framework could accelerate biomarker discovery for conditions where large patient cohorts are unavailable, such as rare diseases or emerging pathogens^99,100^. For infectious diseases, the approach could identify host response signatures that complement or surpass pathogen-specific tests, particularly valuable when pathogens are difficult to detect directly (e.g., *Salmonella typhi*, causative agent of typhoid fever, is present at <1 cfu/ml of blood during active disease^101^) or when distinguishing active disease from latent infection, colonization, and diseases that are symptomatically/immunologically similar to many other diseases^102^ (e.g., Lyme disease^22^).

The multipurpose utility demonstrated by MAS_TB_META5, meeting WHO performance criteria for prognostic, and screening applications, highlights another advantage of biologically-grounded biomarkers. Our findings show that because the five genes in MAS_TB_META5 mechanistically captured fundamental aspects of host responses and disease biology rather than circumstantial correlations, the signature maintained performance across different disease stages and contexts^103,104^. In fact, MAS_TB_META5 was the top ranked signature based on a composite score computed as the sum of wAUROC, overall success rate of achieving WHO minimal TPP for high sensitivity screening and prognostic tests across all validation cohorts, and the number of different contexts in which WHO TPPs were achieved (**Fig. 6**). Notably, only four signatures in addition to MAS_TB_META5 (viz., Gong_OD_4, Hoang_OD_3, Kaul_3, and MAS_TB_OD4) achieved WHO TPPs across the four contexts. MAS_TB_META5 had higher wAUROC and overall success rate in achieving WHO TPP across all independent cohorts (0.852 and 63.2%, respectively) as compared to the next best performing signature (Gong_OD_4; 0.837 and 61.9%). A noteworthy observation was that Gong_OD_4 was also discovered with a network-based approach for gene selection^62^. While Hoang_OD_3, Kaul_3, and MAS_TB_OD4 achieved WHO TPP for a prognostic test to predict TB progression in a single validation cohort, MAS_TB_META5 achieved the minimum performance metrics in three out of the four cohorts. Notably, MAS_TB_OD4 was the only signature that achieved WHO TPP for a prognostic test for the 24-month window in the GSE79362 cohort (**Fig. 6**). Another notable finding from this analysis was that MAS_TB3a topped the list of five signatures, including LauxdaCosta_OD_3, Roe_3, Hoang_OD_3, and Kaul_3, that achieved WHO TPP for a high-specificity screening test, i.e., a rule out test, across three relevant contexts (**Fig. S7**). This versatility of monocyte-amplified signatures, demonstrated by the multipurpose utility of the five MAS signatures discovered in this study, could be transformational for clinical practice by enabling single tests that serve multiple purposes: screening high-risk populations, confirming suspected cases, and monitoring treatment response.

Several limitations merit consideration. First, while monocyteMINER was constructed from a diverse multi-ethnic cohort, expanding the approach with additional populations could enhance its completeness. Second, although we demonstrated the approach with atherosclerosis and TB, systematic validation across a broader spectrum of diseases would strengthen claims of generalizability. Third, the current network captures steady-state transcriptional relationships; incorporating dynamic responses to stimuli could reveal additional disease-relevant modules.

Future directions should explore several promising avenues. Integration of single-cell transcriptomics could refine our understanding of monocyte heterogeneity and subset-specific responses to disease^14^. Combining monocyte TRN with those from other immune cells (e.g., neutrophils) could create a multi-dimensional view of immune surveillance, expanding the utility of this approach to a range of chronic and acute diseases^105,106^. The short median life span of circulating monocytes (1-3 days^107^) suggests that MAS signatures are a reflection of recent health history of the patient, which by extension also suggests that the same biomarkers could also predict treatment response. The context-specificity of monocyte responses^108^ and the composition of training and test datasets should enable the discovery of biomarkers that can distinguish across disease stages and between diseases with overlapping clinical presentations, as we have demonstrated with the ability of MAS_TB_META5 to distinguish aTB from other respiratory illnesses in symptomatic patients. Finally, the biological interpretability of network-derived signatures for atherosclerosis and TB illustrate how monocyteMINER-based approach could uncover actionable biomarkers that can guide development of targeted therapeutics by identifying key regulatory nodes driving pathological responses.

The success of MAS_TB_META5 in meeting WHO TPP across multiple applications demonstrates that the path to clinically useful biomarkers lies not in mining ever-larger datasets with more powerful algorithms, but in leveraging and understanding the fundamental biology of how immune cells encode information about disease. By treating monocytes as biological sensors and their transcriptional networks as the decoding machinery, we can discover biomarkers that are not only statistically robust but mechanistically meaningful. This approach promises to accelerate precision medicine by providing clinicians with diagnostic tools that reflect the actual biological processes driving disease, enabling earlier detection, more accurate diagnosis, and ultimately better patient outcomes.

## METHODS

### Inference of a transcriptional regulatory network model for human monocytes

A publicly available transcriptional compendium of 1,202 donor-isolated monocyte samples from the Multi-Ethnic Study of Atherosclerosis^32^ (Gene Expression Omnibus—GEO accession ID GSE56045) was used together with transcription factor (TF) binding profile information previously reported on the TF binding site database (TFBSDB)^109^ to infer a modular model of the transcriptional network of human monocytes using the Mechanistic Inference of Node Edge Relationships (MINER) algorithm^31^. Briefly, MINER organizes co-expressed genes into regulons (i.e., clusters of co-regulated genes) based on evidence of their co-regulation by specific TFs (derived from TFBSDB) across transcriptome subsets. Next, activity of each regulon is quantized into over, under, or neutral based on distribution of member gene expression patterns across the transcriptional compendium (**Fig. S1A**). Finally, regulons with similar activity profiles across samples are clustered into transcriptional “programs” to generate the full network model (monocyteMINER; **Fig. S1B**), which is then used to uncover transcriptional networks that differentiate groups of interest (e.g., active TB cases vs healthy controls). Atherosclerosis network discovery and CAD/AMI validation Methods are described in the **Extended Data**.

### Discovery of monocyte-amplified gene signatures for aTB

Monocyte transcriptomes of four healthy individuals and seven aTB patients were downloaded from GEO (accession ID: GSE19943^35^), and whole blood transcriptomes of 54 individuals diagnosed with aTB and 127 individuals diagnosed with OD from the ArrayExpress repository (accession ID E-MTAB-8290^36^). Raw intensities of the 11 monocyte transcriptomes were background corrected (with the *normexp* method), and quantile normalized across arrays (with the *quantile* method) using the Limma R package^110^. The normalized data was summarized at the RefSeq ID level by averaging the signal of probes associated with the same RefSeq ID (based on the annotation of the GPL6947 microarray platform). Differential expression analysis was performed using a Bayesian T-test, as explained above. Genes with raw p-value < 0.01 and absolute log2 fold-change >1 were considered DEGs. These cutoffs identified 110 DEGs, significantly reproducing previous findings from analysis of whole blood transcriptomes from the same TB cohort^35^. For the second dataset with aTB and OD cases, we performed differential expression analysis using DESeq2 and used adjusted p-value < 0.05 and absolute log2 fold-change > 1 to identify 175 DEGs. monocyteMINER regulons enriched with the DEGs were identified based on a hypergeometric test (p-value < 0.05). Altogether 17 and 21 regulons were significantly enriched with the DEGs from the aTB vs. OD, and aTB vs. healthy/LTBI contexts, respectively. DEGs within these 17 regulons and 21 regulons were independently aggregated into candidate 11- and 27-gene signatures, respectively, for differentiating aTB from OD and healthy/LTBI cases, respectively.

### Function enrichment analysis of monocyteMINER regulons

Function enrichment of regulons was performed using ontology, hallmark, and cell type signature gene sets from the Human Molecular Signatures database (MSigDB)^67^ implemented in the *msigdbr* R package^111^. Only terms with raw p-value and adj. p-value < 0.05, q-value < 0.2, and associated with at least five (and no more than 500) genes were considered enriched.

### Definition of minimal gene signatures for detecting aTB

The 11-gene signature for aTB vs OD was reduced further by evaluating the performance of all 550 distinct two-, three- and four-gene subsets on detecting aTB in the original discovery cohort from Turner et al., and a second independent whole blood transcriptomes dataset of 64 OD patients and 35 culture-confirmed aTB patients from GSE39939^45^, which was processed using methods described earlier for microarray-based transcriptome datasets. Normalized data was summarized at the gene level by computing the median intensity across all probes associated with the same gene (based on the annotation of the GPL10558 microarray platform). A disease score computed as the algorithmic mean of the normalized expression values of member genes within each reduced gene signature (all genes were upregulated in aTB) was used to evaluate performance of detecting aTB cases in each dataset. AUROCs across 1,000 bootstraps of each dataset was computed for all 550 candidate gene signatures. In each bootstrap, 50% of the aTB patients and OD patients were randomly selected and their TB scores calculated to maintain the original proportion of aTB to OD cases. As a control, we evaluated performance of the initial 11-gene signature in the same set of random bootstraps. Additionally, the performance of signatures comprised of co-regulated genes within each of 17 DEG-enriched monocyteMINER regulons was also individually evaluated. Finally, candidate signatures were ranked using an approach similar to the concept of weighted AUROC^44^ to account for differences in cohort size across datasets. Two minimal gene signatures for aTB were selected for downstream analyses: MAS_TB_OD4, the best AUROC-based ranked gene set among all two-, three- and four-gene signatures, and MAS_TB_OD3, the 3-gene signature associated with regulon # 5050, which was the best ranked regulon-based signature with less than five DEGs (**Table 2**). A five gene threshold was set to facilitate future clinical translation while comparing performance with previously reported best-performing signatures (such as Sweeney3, Hoang3, and Roe3).

For aTB vs HC/LTBI, the initial 27-gene signature was refined into a 17-gene signature (MAS_TB17) by comparing the eigengenes of all monocyteMINER regulons between the aTB and the control group in the discovery cohort (i.e., monocyte transcriptomes). The 17-gene signature was comprised of DEGs within regulons (#3664 and #3685) deemed differentially active based on both DEG enrichment and eigengene comparison with a relaxed adjusted t-test p-value cutoff of 0.29. Similar to the approach described for the discovery of candidate gene signatures for aTB vs OD, we evaluated the performance of 816 two- and three-gene signatures from MAS_TB17, 21 signatures derived from co-regulated gene sets within each of the 21 DEG-enriched monocyteMINER regulons, and MAS_TB17. Performance was evaluated with 1,000 bootstraps as explained above using an independent compendium of whole blood transcriptomes consisting of 54 individuals with aTB, 69 individuals with LTBI, and 24 uninfected (“healthy”) individuals^35^. This analysis identified MAS_TB3a, the 3-gene signature with the highest average AUROC, and MAS_TB3b, the best performing signature among the regulon-based signatures, as two candidate gene signatures for aTB vs healthy/LTBI for downstream analyses (**Table 2**).

### Validation cohorts for MAS TB signatures

Performance of gene signatures was evaluated using datasets from independent cohorts from clinically relevant settings where TB detection is urgently needed (**Table 1**). We compiled publicly available blood transcriptomes from cohorts that include aTB cases, healthy individuals (with and without latent TB infection), ODs with similar symptomatology, healthy individuals that progress into aTB, healthy individuals that do not develop active disease, and advanced HIV patients with and without aTB. We used the work of Warsinske et al.^44^ as an initial survey of relevant datasets that we further complemented with additional literature review^5,39^. Representative cohorts with ≥ 30 total samples (with the exception of the GSE162164 dataset that was combined with GSE107104—more details below) among the shortlisted datasets were selected to evaluate performance of the MAS_TB signatures across different geographic locations, sequencing platforms, control groups (e.g., Mtb uninfected, LTBI, OD), clinical forms of TB (pediatric and adult pulmonary TB, and extrapulmonary TB) and HIV status. In the context of aTB vs OD, we only considered cohorts in which the OD group was not limited to a pre-selected set of pulmonary diseases to focus instead on datasets that emulate observational studies in which patients with TB and a wide range of non-TB diseases are enrolled when seeking medical care^36^. In summary, we used 18 validation cohorts, including a qRT-PCR dataset, nine RNAseq datasets, and eight microarray-based datasets (**Table 1**), to rigorously evaluate and compare the performance of MAS_TB signatures with previously reported TB signatures. Notably, performance of each signature datasets was evaluated on independent datasets, excluding datasets used for its discovery. For example, the Turner et al. dataset and the GSE39939 dataset (for aTB vs OD), were used for the discovery of MAS_OD3 and MAS_OD4, and therefore excluded from performance assessments of these two signatures and MAS_TB_META5.

### Performance assessment

We used the PLAGE score in the TBSignatureProfiler R package^37^, which was demonstrated previously to outperform other scoring metrics^39^, to estimate and compare performance of the MAS_TB signatures and 77 previously reported TB signatures. Raw intensities of microarray datasets (GSE39939, GSE39940, GSE28623, GSE37250, GSE56153) were downloaded from GEO and normalized with background correction, followed by quantile normalization, and finally gene level summary by computing the median signal intensity among relevant probes^80^. Relative to the original Agilent gProcessedSignal, the normalization workflow yielded significantly better average performance results for all 77 signatures on the GSE28623 dataset (t-test p-value < 0.05), which was generated with an Agilent array. For the GSE144127 dataset^46^, we used the original normalized expression values for comparative purposes. We confirmed with a t-test that our normalization workflow did not significantly change the distribution of AUROC values for all gene signatures (p-value > 0.7). For the remaining two validation microarray-based datasets, GSE73408 and GSE62525, we used the original authors’ normalized values. GSE73408 was normalized by Walter et al. using the recommended normalization method for Affymetrix arrays, robust multi-array average^112^. For GSE62525, generated with a Phalanx OneArray platform, Lee et al. used the manufacturer-recommended data processing software^41^.

There were nine RNAseq datasets among the validation cohorts. When possible, we followed TBSignatureProfiler recommendations and used log_2_(counts per million) (log_2_CPM) transformed values as input for performance assessments. For six RNAseq datasets (GSE79362, GSE84076, GSE94438, GSE152218, GSE162164 and the Turner et al. dataset), we downloaded raw read counts from GEO (or ArrayExpress) and then applied a log_2_CPM transformation. For the GSE107104 dataset, we used the raw counts preloaded in the TBSignatureProfiler R package and applied a log_2_CPM transformation. For the GSE79362 dataset, we analyzed two versions of the data. In the first version, we used NCBI-generated RNA-seq raw count data available in GEO. The second version was generated by downloading and using as input the splice junction counts estimated by Zak et al^65^. In the latter, the downloaded expression matrix was summarized at the gene level by computing the median across the log2CPM transformed counts of all junctions linked to each gene. Similarly, for GSE84076, we used NCBI-generated RNA-seq count data. We confirmed that the AUROC distribution reproduced results generated using normalized data from the original publication^113^. To allow comparative assessments with the original studies, we used DESeq2 normalization for GSE101205 and GSE112104 datasets, which is compatible with TBSignatureProfiler. For the qRT-PCR dataset reported by Mendelsohn et al.^56^, we used as input for our scoring scheme delta Ct values (see below). Due to discrepancies in gene names across platforms and versions of the human genome annotation, for each validation dataset we checked the coverage of all candidate TB signatures, manually curating gene synonyms to reduce missing data for signatures with four or less genes. We limited this manual curation to small gene sets due to the demanding nature of this task, the expectation that smaller gene signatures would be the ones most affected by the incomplete coverage, and the recognition of this potential caveat in previous work. Importantly, for GSE39939 and GSE144127, which contain culture-confirmed and unconfirmed TB cases, analyses shown in the main text were performed using the culture-confirmed TB cases. Results of analysis on the full datasets are in supplement.

For advanced HIV and TB co-infection, we combined the results of two small validation cohorts, GSE107104 and GSE162164, to increase the statistical power of the analysis. For each dataset, we first computed PLAGE scores using log2CPM values. Then, we combined the PLAGE scores of both datasets with the *ComBat* batch-correction function^114^ in the SVA R package^115^, incorporating TB status as a covariate. We tested two additional methods to integrate the two datasets. Results in **Fig. 4A-C** were generated using the integration approach described above because it resulted in the highest AUROC value for Kulkarni2 (the only signature discovered using these datasets), had the highest number (8 out of 34) of previous TB signatures achieving the TPP for a high-sensitivity screening test (vs. five and six for the alternative approaches), had Pearson correlation > 0.996 with PLAGE scores computed in each individual dataset for all TB signatures of interest (i.e., with 10 genes or less), and had higher AUROC than the other two alternatives for ∼69% of previously proposed TB signatures. In a second approach, PLAGE scores for each dataset were independently z-scored and then merged. This approach of integration was feasible because z-scores can be compared across datasets, and the two datasets in question were similar with respect to distributions of z-scores (**Fig. S5**) and the ratios of active TB to control (i.e., advanced HIV only) cases. In a third approach described by Clarke et al.^22^, log2 transformed raw counts in each dataset were quantile normalized, z-scored, combined using batch-correction with *ComBat*, and then used to compute PLAGE scores with TBSignatureProfiler.

Performance of signatures in predicting TB progression was evaluated against established WHO guidelines for a progression test, using datasets from four cohorts (four countries: Brazil, S. Africa, The Gambia, and Ethiopia) across multiple time intervals (from six months up to five years) between sampling and TB diagnosis (**Table 1**). In the multi-country GSE94438 dataset^57^, there were 64 TB progressor transcriptomes and 208 non-progressors transcriptomes collected at baseline (i.e., enrollment after exposure) in the 2-year interval study. In the 6-month interval window analysis, we used the compendium-associated public metadata to reconstruct timelines and select the pre-disease transcriptome collected in the six-month window before confirmed TB diagnosis (as previously done by Warsinske et al.^116^). This resulted in 47 TB progressor transcriptomes (∼76% of cases with time to disease ≥ 4 months) and 259 non-progressor transcriptomes (only the transcriptome collected at the first time point of each non-progressor was used to avoid redundancy). Similar analyses were performed for the 12-month and 24-month intervals, and by country. For each sample-to-disease interval, we selected the transcriptomes that preceded disease by no more than the target time interval. When multiple transcriptomes were collected within the desired time interval for a given patient, the transcriptome collected at the largest time interval between sampling and time of confirmed disease diagnosis was selected. This workflow simulates a real-world situation when people may not be immediately tested after TB exposure (or diagnosis of an index case) and discovered to have been exposed to the disease at a later time. Additionally, this approach efficiently leverages the longitudinal nature of the study to maximize the number of TB progression cases that can be analyzed (e.g., there were only 24 participants that developed TB in the next 6 months after enrollment but 47 that could be used in our analysis). As in the original study, samples of patients that progressed towards aTB at least three months after enrollment were included in the analysis.

The Brazilian cohort A for TB progression^56^ included qRT-PCR profiles for just three out of the five genes (ANKRD22, VAMP5 and GBP5) in MAS_TB_META5. Hence, performance of a partial gene signature for MAS_TB_META5 was evaluated for the two-year interval (using blood collected at enrollment), and for 6-month, 9-month and 12-month intervals from enrollment (as performed by Mendelsohn et al. for previous TB signatures). Mendelsohn et al. implemented three scoring schemes for each signature: (i) normalized Ct values, (ii) batch-corrected scores, and (iii) normalized scores with log- and z-score transformation^56^. Consistent with findings reported by Muwanga et al.^59^, we did not observe a clear benefit of the two latter schemes that utilized batch-correction and additional transformations. Thus, we simply scored each sample for MAS_TB_META5 as the sum of delta Ct values for ANKRD22, VAMP5 and GBP5. Similar to Rajan5, one of the three signatures that achieved WHO TPP for TB progression for the < 9-month interval^56^, MAS_TB_META5 performed significantly better without batch-correction, thus we used MAS_TB_META5 raw score for downstream analyses.

GSE79362 was generated from a cohort of South African adolescents (ages 12-18 years old) followed up to 24 months for TB progression^65^. We included in our analysis transcriptomes from the initial blood draw (baseline) of participants assigned to the “training set” defined in the study. While inspecting the metadata associated with this dataset, we were unable to identify the baseline transcriptomes of participants assigned to the “test set”. Ten out of the 33 TB progressors did not include a transcriptome profile for day zero (for eight TB progressors, the first sample was collected on day 180, and for two participants the first sample was collected on day 360). For this reason, we used the baseline transcriptomes of all individuals in the training set (representing a target window up to 24 months). While this effectively limited the number of samples for both TB progressors and non-progressors to a single transcriptome, the late enrollment of participants models a real-world scenario. Further, due to challenges in converting junction read counts (used by Zak et al.) into gene level read counts (the standard input for TBSignatureProfiler), the performance assessment of all signatures was evaluated using raw read counts in the GSE79362 dataset.

In the GSE112104 dataset (Brazilian Cohort B), five participants diagnosed with aTB in the first two months post-enrollment were excluded from the analysis, as done in the original study^66^. This reduced the total number of TB progressors in this study to 11.

### Identification of top tier performing TB signatures

A top tier performing signature on a given cohort was defined as a signature with performance (i.e., AUROC) that was indistinguishable relative to that of the best performing signature on the same cohort^5^. Significance of difference between AUROCs was determined using the *roc.test* function in the pROC R package^117^. Only differences with p-value < 0.05 were considered significant. Pairwise comparisons were performed using the Delong method when possible. For other cases, bootstrapping based comparison was used^118^.

## AUTHOR CONTRIBUTIONS

Conceptualization, M.L.A.-O. and N.S.B.; Data curation, M.L.A.-O.; Formal Analysis, M.L.A.-O.; Funding acquisition, M.L.A.-O and N.S.B.; Investigation, M.L.A.-O. and N.S.B.; Methodology, M.L.A.-O. and W.-J.W.; Project administration, M.L.A.-O. and N.S.B.; Resources, M.L.A.-O. and N.S.B.; Software, M.L.A.-O. and W.-J.W.; Supervision, N.S.B.; Visualization, M.L.A.-O. and N.S.B.; Writing, M.L.A.-O. and N.S.B.

## Supporting information

Data Set 2

Data Set 3

Data Set 1

## ACKNOWLEDGEMENTS

We thank current and past members of the Baliga lab for critical discussions and feedback. We especially thank Dr. Serdar Turkarslan, Dr. James Park, and Dr. Carl Murie for their input on MINER use. We thank Dr. Jacob Valenzuela for his feedback. We also thank Dr. Thomas R. Hawn and Dr. Thuong Nguyen Thuy Thuong for their feedback. This work was supported by the National Institutes of Health grants 5R01AI128215-09 and 5R01AI141953-05 (N.S.B.), the Gates Foundation INV-092174 (N.S.B.). This work was also supported by the Tuberculosis Research Unit grant U19AI162583 (Jeffery S. Cox at UC Berkeley and Thomas R. Hawn at University of Washington).

## CODE AVAILABILITY

All scripts used in the analyses described in this work can be found at https://github.com/marioluisao/Monocyte-Amplified-Transcriptional-Signatures-of-Human-Diseases.

## SOFTWARE

Python 3.12.1 was used to generate the monocyteMINER model with the MINER algorithm. All computational analyses of the monocyteMINER model, biomarker discovery and gene signature validation were performed on R version 4.4.1^119^. Networks visualization was generated with Cytoscape 3.10.0^120^. Inkscape was used to generate composite figures.

## REFERENCES

1. Ren, A. H., Fiala, C. A., Diamandis, E. P. & Kulasingam, V. Pitfalls in cancer biomarker discovery and validation with emphasis on circulating tumor DNA. *Cancer Epidemiology*, Biomarkers & Prevention 29, 2568–2574 (2020).

2. T Dudley, J. & Butte, A. J. Identification of discriminating biomarkers for human disease using integrative network biology. in Pacific Symposium on Biocomputing. Pacific Symposium on Biocomputing 27 (2009).

3. Frangogiannis, N. G. Biomarkers: hopes and challenges in the path from discovery to clinical practice. Translational Research 159, 197–204 (2012).

4. Whitcomb, B. W. & Schisterman, E. F. Assays with lower detection limits: implications for epidemiological investigations. Paediatric and perinatal epidemiology 22, 597–602 (2008).

5. Gupta, R. K. et al. Concise whole blood transcriptional signatures for incipient tuberculosis: a systematic review and patient-level pooled meta-analysis. The Lancet Respiratory Medicine 8, 395–406 (2020).

6. Magis, A. T. et al. Untargeted longitudinal analysis of a wellness cohort identifies markers of metastatic cancer years prior to diagnosis. Scientific reports 10, 16275 (2020).

7. Bordeianu, G. et al. Circulating Biomarkers for Laboratory Diagnostics of Atherosclerosis—Literature Review. Diagnostics 12, 3141 (2022).

8. Pedrotty, D. M., Morley, M. P. & Cappola, T. P. Transcriptomic biomarkers of cardiovascular disease. Progress in cardiovascular diseases 55, 64–69 (2012).

9. Levy, H. et al. Transcriptional signatures as a disease-specific and predictive inflammatory biomarker for type 1 diabetes. Genes & Immunity 13, 593–604 (2012).

10. Lake, J., Storm, C. S., Makarious, M. B. & Bandres-Ciga, S. Genetic and transcriptomic biomarkers in neurodegenerative diseases: current situation and the road ahead. Cells 10, 1030 (2021).

11. Esmail, H., Cobelens, F. & Goletti, D. Transcriptional biomarkers for predicting development of TB: progress and clinical considerations. European Respiratory Journal (2020).

12. Ng, S., Masarone, S., Watson, D. & Barnes, M. R. The benefits and pitfalls of machine learning for biomarker discovery. Cell and tissue research 394, 17–31 (2023).

13. Maher, A. K. et al. Transcriptional reprogramming from innate immune functions to a pro-thrombotic signature by monocytes in COVID-19. Nature Communications 13, 7947 (2022).

14. Hillman, H. et al. Single-cell profiling reveals distinct subsets of CD14+ monocytes drive blood immune signatures of active tuberculosis. Frontiers in immunology 13, 1087010 (2023).

15. Schlachetzki, J. C. et al. A monocyte gene expression signature in the early clinical course of Parkinson’s disease. Scientific Reports 8, 10757 (2018).

16. Vallania, F. et al. Multicohort analysis identifies monocyte gene signatures to accurately monitor subset-specific changes in human diseases. Frontiers in Immunology 12, 659255 (2021).

17. Cassetta, L. et al. Human tumor-associated macrophage and monocyte transcriptional landscapes reveal cancer-specific reprogramming, biomarkers, and therapeutic targets. Cancer cell 35, 588–602 (2019).

18. Epelman, S., Lavine, K. J. & Randolph, G. J. Origin and functions of tissue macrophages. Immunity 41, 21–35 (2014).

19. Rodero, M. P. et al. Immune surveillance of the lung by migrating tissue monocytes. Elife 4, e07847 (2015).

20. Gerhardt, T. & Ley, K. Monocyte trafficking across the vessel wall. Cardiovascular research 107, 321–330 (2015).

21. Singhania, A. et al. A modular transcriptional signature identifies phenotypic heterogeneity of human tuberculosis infection. Nature communications 9, 2308 (2018).

22. Clarke, D. J. et al. Predicting lyme disease from patients’ peripheral blood mononuclear cells profiled with RNA-sequencing. Frontiers in immunology 12, 636289 (2021).

23. Feng, A. et al. CD16+ monocytes in breast cancer patients: expanded by monocyte chemoattractant protein-1 and may be useful for early diagnosis. Clinical & Experimental Immunology 164, 57–65 (2011).

24. World Health Organization. Global Tuberculosis Report 2024. https://www.who.int/teams/global-programme-on-tuberculosis-and-lung-health/tb-reports/global-tuberculosis-report-2024.

25. Pai, M., Dewan, P. K. & Swaminathan, S. Transforming tuberculosis diagnosis. Nature Microbiology 8, 756–759 (2023).

26. World Health Organization. High Priority Target Product Profiles for New Tuberculosis Diagnostics: Report of a Consensus Meeting, 28-29 April 2014, Geneva, Switzerland. (2014).

27. Gill, C. M., Dolan, L., Piggott, L. M. & McLaughlin, A. M. New developments in tuberculosis diagnosis and treatment. Breathe 18, (2022).

28. Diel, R. et al. Interferon-γ release assays for the diagnosis of latent Mycobacterium tuberculosis infection: a systematic review and meta-analysis. European Respiratory Journal 37, 88–99 (2010).

29. Mendelsohn, S. C. et al. Prospective multicentre head-to-head validation of host blood transcriptomic biomarkers for pulmonary tuberculosis by real-time PCR. Communications medicine 2, 26 (2022).

30. Chakravorty, S. et al. The new Xpert MTB/RIF Ultra: improving detection of Mycobacterium tuberculosis and resistance to rifampin in an assay suitable for point-of-care testing. MBio 8, 10–1128 (2017).

31. Wall, M. A. et al. Genetic program activity delineates risk, relapse, and therapy responsiveness in multiple myeloma. npj Precision Oncology 5, 60 (2021).

32. Liu, Y. et al. Blood monocyte transcriptome and epigenome analyses reveal loci associated with human atherosclerosis. Nature communications 8, 393 (2017).

33. Andreini, D. et al. Whole-blood transcriptional profiles enable early prediction of the presence of coronary atherosclerosis and high-risk plaque features at coronary CT angiography. Biomedicines 10, 1309 (2022).

34. Muse, E. D. et al. A whole blood molecular signature for acute myocardial infarction. Scientific reports 7, 12268 (2017).

35. Berry, M. P. et al. An interferon-inducible neutrophil-driven blood transcriptional signature in human tuberculosis. Nature 466, 973–977 (2010).

36. Turner, C. T. et al. Blood transcriptional biomarkers for active pulmonary tuberculosis in a high-burden setting: a prospective, observational, diagnostic accuracy study. The Lancet Respiratory Medicine 8, 407–419 (2020).

37. Johnson, W. E. et al. Comparing tuberculosis gene signatures in malnourished individuals using the TBSignatureProfiler. BMC infectious diseases 21, 1–13 (2021).

38. Tomfohr, J., Lu, J. & Kepler, T. B. Pathway level analysis of gene expression using singular value decomposition. BMC bioinformatics 6, 1–11 (2005).

39. Wang, X. et al. Comparison of gene set scoring methods for reproducible evaluation of tuberculosis gene signatures. BMC Infectious Diseases 24, 610 (2024).

40. Sambarey, A. et al. Unbiased identification of blood-based biomarkers for pulmonary tuberculosis by modeling and mining molecular interaction networks. EBioMedicine 15, 112–126 (2017).

41. Lee, S.-W. et al. Gene expression profiling identifies candidate biomarkers for active and latent tuberculosis. BMC bioinformatics 17, S3 (2016).

42. Verhagen, L. M. et al. A predictive signature gene set for discriminating active from latent tuberculosis in Warao Amerindian children. BMC genomics 14, 74 (2013).

43. Leong, S. et al. Existing blood transcriptional classifiers accurately discriminate active tuberculosis from latent infection in individuals from south India. Tuberculosis 109, 41–51 (2018).

44. Warsinske, H., Vashisht, R. & Khatri, P. Host-response-based gene signatures for tuberculosis diagnosis: A systematic comparison of 16 signatures. PLoS medicine 16, e1002786 (2019).

45. Anderson, S. T. et al. Diagnosis of childhood tuberculosis and host RNA expression in Africa. New England Journal of Medicine 370, 1712–1723 (2014).

46. Hoang, L. T. et al. Transcriptomic signatures for diagnosing tuberculosis in clinical practice: a prospective, multicentre cohort study. The Lancet Infectious Diseases 21, 366–375 (2021).

47. Chang, A. et al. Circulating cell-free RNA in blood as a host response biomarker for detection of tuberculosis. Nature Communications 15, 4949 (2024).

48. Müller, M. et al. Immune reconstitution inflammatory syndrome in patients starting antiretroviral therapy for HIV infection: a systematic review and meta-analysis. The Lancet infectious diseases 10, 251–261 (2010).

49. Walker, N. F., Scriven, J., Meintjes, G. & Wilkinson, R. J. Immune reconstitution inflammatory syndrome in HIV-infected patients. HIV/AIDS-Research and Palliative Care 49–64 (2015).

50. Mann, T. et al. Blood RNA biomarkers for tuberculosis screening in people living with HIV before antiretroviral therapy initiation: a diagnostic accuracy study. The Lancet Global Health 12, e783–e792 (2024).

51. Antinori, A. et al. Late presentation of HIV infection: a consensus definition. HIV medicine 12, 61–64 (2011).

52. Gupta, R. K., Lucas, S. B., Fielding, K. L. & Lawn, S. D. Prevalence of tuberculosis in post-mortem studies of HIV-infected adults and children in resource-limited settings: a systematic review and meta-analysis. Aids 29, 1987–2002 (2015).

53. Kulkarni, V. et al. A two-gene signature for tuberculosis diagnosis in persons with advanced HIV. Frontiers in immunology 12, 631165 (2021).

54. Verma, S. et al. Tuberculosis in advanced HIV infection is associated with increased expression of IFNγ and its downstream targets. BMC infectious diseases 18, 1–13 (2018).

55. Penn-Nicholson, A. et al. RISK6, a 6-gene transcriptomic signature of TB disease risk, diagnosis and treatment response. Scientific reports 10, 8629 (2020).

56. Mendelsohn, S. C. et al. Transcriptomic Signatures of Progression to Tuberculosis Disease Among Close Contacts in Brazil. The Journal of Infectious Diseases jiae237 (2024).

57. Suliman, S. et al. Four-gene pan-African blood signature predicts progression to tuberculosis. American journal of respiratory and critical care medicine 197, 1198–1208 (2018).

58. Hamada, Y. et al. Are mRNA based transcriptomic signatures ready for diagnosing tuberculosis in the clinic?-A review of evidence and the technological landscape. EBioMedicine 82, (2022).

59. Muwanga, V. M. et al. Blood transcriptomic signatures for symptomatic tuberculosis in an African multicohort study. European Respiratory Journal 64, (2024).

60. Gliddon, H. D. et al. Identification of reduced host transcriptomic signatures for tuberculosis disease and digital PCR-based validation and quantification. Frontiers in immunology 12, 637164 (2021).

61. Jacobsen, M. et al. Candidate biomarkers for discrimination between infection and disease caused by Mycobacterium tuberculosis. Journal of molecular medicine 85, 613–621 (2007).

62. Gong, Z. et al. The evaluation and validation of blood-derived novel biomarkers for precise and rapid diagnosis of tuberculosis in areas with high-TB burden. Frontiers in Microbiology 12, 650567 (2021).

63. Rajan, J. V. et al. A novel, 5-transcript, whole-blood gene-expression signature for tuberculosis screening among people living with human immunodeficiency virus. Clinical Infectious Diseases 69, 77–83 (2019).

64. Duffy, F. J., Thompson, E. G., Scriba, T. J. & Zak, D. E. Multinomial modelling of TB/HIV co-infection yields a robust predictive signature and generates hypotheses about the HIV+ TB+ disease state. PLoS One 14, e0219322 (2019).

65. Zak, D. E. et al. A blood RNA signature for tuberculosis disease risk: a prospective cohort study. The Lancet 387, 2312–2322 (2016).

66. Leong, S. et al. Cross-validation of existing signatures and derivation of a novel 29-gene transcriptomic signature predictive of progression to TB in a Brazilian cohort of household contacts of pulmonary TB. Tuberculosis 120, 101898 (2020).

67. Liberzon, A. et al. Molecular signatures database (MSigDB) 3.0. Bioinformatics 27, 1739– 1740 (2011).

68. Mahasirimongkol, S. et al. Genome-wide association studies of tuberculosis in Asians identify distinct at-risk locus for young tuberculosis. Journal of human genetics 57, 363–367 (2012).

69. Chen, C. et al. A rare variant at 11p13 is associated with tuberculosis susceptibility in the Han Chinese population. Scientific reports 6, 24016 (2016).

70. Kelly, L. M., Englmeier, U., Lafon, I., Sieweke, M. H. & Graf, T. MafB is an inducer of monocytic differentiation. The EMBO journal (2000).

71. Hikichi, H., Seto, S., Wakabayashi, K., Hijikata, M. & Keicho, N. Transcription factor MAFB controls type I and II interferon response-mediated host immunity in Mycobacterium tuberculosis-infected macrophages. Frontiers in Microbiology 13, 962306 (2022).

72. Sugawara, I., Yamada, H. & Mizuno, S. Relative importance of STAT4 in murine tuberculosis. Journal of medical microbiology 52, 29–34 (2003).

73. Matsui, T., Sakamaki, Y., Hiragi, S. & Fukuda, M. VAMP5 and distinct sets of cognate Q-SNAREs mediate exosome release. Cell Structure and Function 48, 187–198 (2023).

74. Sakurai, C., Yamashita, N., Azuma, K. & Hatsuzawa, K. VAMP5 promotes Fcγ receptor-mediated phagocytosis and regulates phagosome maturation in macrophages. Molecular Biology of the Cell 35, ar44 (2024).

75. Pauwels, A.-M., Trost, M., Beyaert, R. & Hoffmann, E. Patterns, receptors, and signals: regulation of phagosome maturation. Trends in immunology 38, 407–422 (2017).

76. Fujiwara, Y. et al. Guanylate-binding protein 5 is a marker of interferon-γ-induced classically activated macrophages. Clinical & translational immunology 5, e111 (2016).

77. Shenoy, A. R. et al. GBP5 promotes NLRP3 inflammasome assembly and immunity in mammals. Science 336, 481–485 (2012).

78. Ehrt, S. et al. Reprogramming of the macrophage transcriptome in response to interferon-γ and Mycobacterium tuberculosis: signaling roles of nitric oxide synthase-2 and phagocyte oxidase. The Journal of experimental medicine 194, 1123–1140 (2001).

79. Lösslein, A. K. & Henneke, P. Macrophage differentiation and metabolic adaptation in mycobacterial infections. Annual Review of Immunology 43, (2025).

80. Sweeney, T. E., Braviak, L., Tato, C. M. & Khatri, P. Genome-wide expression for diagnosis of pulmonary tuberculosis: a multicohort analysis. The Lancet Respiratory Medicine 4, 213–224 (2016).

81. Martinez, F. O. et al. Genetic programs expressed in resting and IL-4 alternatively activated mouse and human macrophages: similarities and differences. *Blood*, The Journal of the American Society of Hematology 121, e57–e69 (2013).

82. Huang, Z. et al. Mycobacterium tuberculosis-induced polarization of human macrophage orchestrates the formation and development of tuberculous granulomas in vitro. PloS one 10, e0129744 (2015).

83. Ahmad, F. et al. Macrophage: a cell with many faces and functions in tuberculosis. Frontiers in immunology 13, 747799 (2022).

84. Palucci, I. et al. Transglutaminase type 2 plays a key role in the pathogenesis of Mycobacterium tuberculosis infection. Journal of internal medicine 283, 303–313 (2018).

85. Palucci, I. et al. Inhibition of transglutaminase 2 as a potential host-directed therapy against Mycobacterium tuberculosis. Frontiers in Immunology 10, 3042 (2020).

86. Utsumi, T. et al. ANKRD22 is an N-myristoylated hairpin-like monotopic membrane protein specifically localized to lipid droplets. Scientific Reports 11, 19233 (2021).

87. Zhang, S., Liu, Y., Zhang, X.-L., Sun, Y. & Lu, Z.-H. ANKRD22 aggravates sepsis-induced ARDS and promotes pulmonary M1 macrophage polarization. Journal of Translational Autoimmunity 8, 100228 (2024).

88. Shan, Q. et al. Distinguish active tuberculosis with an immune-related signature and molecule subtypes: a multi-cohort analysis. Scientific Reports 14, 29564 (2024).

89. Briken, V., Ahlbrand, S. E. & Shah, S. Mycobacterium tuberculosis and the host cell inflammasome: a complex relationship. Frontiers in cellular and infection microbiology 3, 62 (2013).

90. Saiga, H. et al. Critical role of AIM2 in Mycobacterium tuberculosis infection. International immunology 24, 637–644 (2012).

91. Theobald, S. J., Müller, T. A., Lange, D., Keck, K. & Rybniker, J. The role of inflammasomes as central inflammatory hubs in Mycobacterium tuberculosis infection. Frontiers in Immunology 15, 1436676 (2024).

92. Shah, S. et al. Cutting edge: Mycobacterium tuberculosis but not nonvirulent mycobacteria inhibits IFN-β and AIM2 inflammasome–dependent IL-1β production via its ESX-1 secretion system. The Journal of Immunology 191, 3514–3518 (2013).

93. Ioannidis, J. P. & Bossuyt, P. M. Waste, leaks, and failures in the biomarker pipeline. Clinical chemistry 63, 963–972 (2017).

94. Rifai, N., Gillette, M. A. & Carr, S. A. Protein biomarker discovery and validation: the long and uncertain path to clinical utility. Nature biotechnology 24, 971–983 (2006).

95. Clarke, R. et al. The properties of high-dimensional data spaces: implications for exploring gene and protein expression data. Nature reviews cancer 8, 37–49 (2008).

96. Chaussabel, D. et al. A modular analysis framework for blood genomics studies: application to systemic lupus erythematosus. Immunity 29, 150–164 (2008).

97. Newman, A. M. et al. Determining cell type abundance and expression from bulk tissues with digital cytometry. Nature biotechnology 37, 773–782 (2019).

98. Shen-Orr, S. S. & Gaujoux, R. Computational deconvolution: extracting cell type-specific information from heterogeneous samples. Current opinion in immunology 25, 571–578 (2013).

99. Simonetta, I., Tuttolomondo, A., Daidone, M. & Pinto, A. Biomarkers in Anderson–Fabry Disease. International Journal of Molecular Sciences 21, 8080 (2020).

100. Bax, B. E. Biomarkers in rare diseases. International journal of molecular sciences vol. 22 673 (2021).

101. Wain, J. et al. Quantitation of bacteria in blood of typhoid fever patients and relationship between counts and clinical features, transmissibility, and antibiotic resistance. Journal of clinical microbiology 36, 1683–1687 (1998).

102. Kobayashi, T., Higgins, Y., Melia, M. T. & Auwaerter, P. G. Mistaken identity: many diagnoses are frequently misattributed to Lyme disease. The American journal of medicine 135, 503–511 (2022).

103. Robinson, W. H., Lindstrom, T. M., Cheung, R. K. & Sokolove, J. Mechanistic biomarkers for clinical decision making in rheumatic diseases. Nature Reviews Rheumatology 9, 267–276 (2013).

104. Atallah, J. & Mansour, M. K. Implications of using host response-based molecular diagnostics on the management of bacterial and viral infections: a review. Frontiers in Medicine 9, 805107 (2022).

105. Patel, A. A., Ginhoux, F. & Yona, S. Monocytes, macrophages, dendritic cells and neutrophils: an update on lifespan kinetics in health and disease. Immunology 163, 250–261 (2021).

106. Ribeiro-Gomes, F. L., Peters, N. C., Debrabant, A. & Sacks, D. L. Efficient capture of infected neutrophils by dendritic cells in the skin inhibits the early anti-leishmania response. PLoS pathogens 8, e1002536 (2012).

107. Whitelaw, D. & Bell, M. The intravascular lifespan of monocytes. Blood 28, 455–464 (1966).

108. Amarasinghe, H. E. et al. Mapping the epigenomic landscape of human monocytes following innate immune activation reveals context-specific mechanisms driving endotoxin tolerance. BMC genomics 24, 595 (2023).

109. Plaisier, C. L. et al. Causal mechanistic regulatory network for glioblastoma deciphered using systems genetics network analysis. Cell systems 3, 172–186 (2016).

110. Ritchie, M. E. et al. limma powers differential expression analyses for RNA-sequencing and microarray studies. Nucleic acids research 43, e47–e47 (2015).

111. Dolgalev, I. Msigdbr: MSigDB Gene Sets for Multiple Organisms in a Tidy Data Format. (2025).

112. Walter, N. D. et al. Blood transcriptional biomarkers for active tuberculosis among patients in the United States: a case-control study with systematic cross-classifier evaluation. Journal of clinical microbiology 54, 274–282 (2016).

113. De Araujo, L. S. et al. Transcriptomic biomarkers for tuberculosis: evaluation of DOCK9. EPHA4, and NPC2 mRNA expression in peripheral blood. Frontiers in microbiology 7, 1586 (2016).

114. Johnson, W. E., Li, C. & Rabinovic, A. Adjusting batch effects in microarray expression data using empirical Bayes methods. Biostatistics 8, 118–127 (2007).

115. Leek, J. T., Johnson, W. E., Parker, H. S., Jaffe, A. E. & Storey, J. D. The sva package for removing batch effects and other unwanted variation in high-throughput experiments. Bioinformatics 28, 882–883 (2012).

116. Warsinske, H. C. et al. Assessment of validity of a blood-based 3-gene signature score for progression and diagnosis of tuberculosis, disease severity, and treatment response. JAMA network open 1, e183779–e183779 (2018).

117. Robin, X. et al. pROC: an open-source package for R and S+ to analyze and compare ROC curves. BMC bioinformatics 12, 77 (2011).

118. DeLong, E. R., DeLong, D. M. & Clarke-Pearson, D. L. Comparing the areas under two or more correlated receiver operating characteristic curves: a nonparametric approach. Biometrics 837–845 (1988).

119. R Core Team. R: A Language and Environment for Statistical Computing. (R Foundation for Statistical Computing, Vienna, Austria, 2024).

120. Shannon, P. et al. Cytoscape: a software environment for integrated models of biomolecular interaction networks. Genome Res 13, 2498–2504 (2003).

121. Kaforou, M. et al. Detection of tuberculosis in HIV-infected and-uninfected African adults using whole blood RNA expression signatures: a case-control study. PLoS medicine 10, e1001538 (2013).

122. Maertzdorf, J. et al. Functional correlations of pathogenesis-driven gene expression signatures in tuberculosis. PloS one 6, e26938 (2011).

123. Ottenhoff, T. H. et al. Genome-wide expression profiling identifies type 1 interferon response pathways in active tuberculosis. (2012).

