## Extended Data for "Monocyte-amplified transcriptional signatures of human diseases"

This document presents Extended Data complementing the main text. Four sections provide full detail on: (1) monocyteMINER network construction; (2) atherosclerosis proof-of-concept results including CAD and AMI biomarker discovery and validation; (3) benchmarking of MAS_TB17, MAS_TB3a, and MAS_TB3b intermediate signatures against prior published signatures; and (4) the full per-signature comparative performance enumeration. Extended Data Online Methods provides the complete atherosclerosis analysis methods omitted from the main text.

#### Extended Data 1 | monocyteMINER Network Construction

We hypothesized that upon encountering disease tissue or their molecular signals (e.g., cytokines), the transcriptional response of monocytes internalizes information that can be decoded using a systems-scale model of a monocyte cell type-specific TRN to detect the disease and its underlying biology. To test this hypothesis, we first constructed a systems scale model of the human monocyte TRN using mechanistic inference of node-edge relationships (MINER)[1] and systems genetics network analysis (SYGNAL)[2]. In brief, we applied MINER to a compendium of monocyte transcriptomes from 1,202 "healthy" individuals from the Multi-Ethnic Study of Atherosclerosis (MESA)[3] to uncover co-regulated gene modules (regulons), based on evidence of co-expression of member genes across sub-populations of subjects, with SYGNAL-uncovered evidence of binding locations for at least one transcription factor (TF) across promoters of all genes within each regulon (**Fig. 1A**). In so doing, MINER generated a systems scale model of the TRN of monocytes (monocyteMINER), wherein 529 TFs (out of the 708 human TFs in the Transcription Factor Binding Site Database (TFBS_DB))[2] were mechanistically implicated in a combinatorial scheme for coordinate regulation of 10,567 genes organized into 5,682 regulons (**Fig. 1A, Fig. S1,** and **Data Set S1**).

#### Extended Data 2 | Atherosclerosis Proof-of-Concept: Perturbed Network and Biomarkers for CAD and AMI

We investigated the feasibility of using the monocyteMINER model to decode mechanistic biology of atherosclerosis from transcriptional responses of monocytes. Through eigengene comparisons (see ***Extended Data Online Methods***), we identified four regulons that were differentially regulated (adjusted p-value < 0.05) between 293 individuals with high-risk for heart disease (i.e., calcium artery coronary (CAC) scan scores > 300[4,5]), and 336 individuals with low risk for heart disease (i.e., CAC scores = 0) from the same cohort (**Fig. 1B**). Two regulons (total 17 genes) were overactive and two regulons (total 14 genes) were underactive in patients with high risk for atherosclerosis. Four TFs (HSF1, IRF9, IKZF1 and DR1) were implicated in direct regulation of these differentially active regulons comprising 31 genes that we will here onwards call the "atherosclerosis-perturbed network" (**Fig. 1C**). At least 22 genes within these four regulons (e.g., CD74, NOD1, and ARAP2) were also coordinately regulated with genes across 32 additional regulons by 14 TFs of which some, including PPARA[6–8], ELK1[9,10], and STAT1[11,12], have been previously associated with atherosclerosis and heart disease (**Fig. 1C**).

Analysis of gene functions and interactions within the atherosclerosis-perturbed network generated an integrated perspective of both previously characterized and novel proinflammatory mechanisms, demonstrating the mechanistic association between this network and atherosclerosis development (**Fig. 1C-D**). Notably, regulon #869 (R869) was significantly overactive in the high-risk group and was enriched for genes associated with positive regulation of interleukin-6 (IL-6) production, such as NOD1, CD74, and TYROBP (**Table S1**). While TYROBP has been reported as a consistently overexpressed gene in plaques of atherosclerosis patients[13], NOD1 upregulation in monocytes is known to promote inflammation[14]. Similarly, CD74 upregulation in atherosclerotic plaques[15] influences monocyte recruitment and migration to lesions due to its interaction with Macrophage Inhibitory Factor (MIF)[16], which is at high concentrations in atherosclerotic lesions[17]. Consequently, CD74 upregulation may result in elevated IL-6 levels in monocyte-derived macrophages through the CD74-MIF interaction[18]. monocyteMINER predicted that R869 was under positive transcriptional control of HSF1, which is consistent with the upregulation of this TF in atherosclerosis animal models[19] and the reduced detection of atherosclerotic lesions in *hsf1* knockout mouse models[20]. Together these findings support the hypothesis that HSF1 drives early atherosclerosis by upregulating R869, which induces IL-6 production and promotes monocyte expansion and differentiation into macrophages

(**Fig. 1D**). This hypothesis aligns with multiple observations: oxidized low-density lipoprotein (oxiLDL) triggers both recruitment of monocytes through proinflammatory cytokine production (including IL-6)[21,22] as well as trained immunity in recruited monocytes[23], while promoting their differentiation into macrophages[24] through a process that requires HSF1[25]. Furthermore, increased HSF1 activity corresponds with the documented upregulation of heat shock proteins (HSPs) in response to oxiLDL[26], consistent with HSF1's established role as an HSP regulator[27].

monocyteMINER-inferred regulatory influences revealed gene interactions between R869 and R4143–the other overactive regulon in patients with elevated CAC scores–that may contribute to the pro-inflammatory state that promotes the development of atherosclerosis[3,28]. Notably, siRNA knockdown of ARID5B, an R4143 gene known to influence IL-6 production in atherosclerosis[3], resulted in reduced expression of R869 genes BEX4, CD74, and TYROBP in THP1 cells[3] (**Fig. 1C-D**). R4143 genes were predicted to be regulated by IRF9, which has been independently associated with neointima formation –a hallmark of both atherosclerosis and injury-induced vascular remodeling[29]. In summary, the atherosclerosis-perturbed network delineated modular organization of genes associated with chronic inflammation and provided novel insights into how their interplay may underlie the accumulation of oxiLDL and promote atherosclerosis development.

High CAC score is a prognostic marker for atherosclerosis[4,5], which in turn leads to the development of a wide range of heart diseases, including CAD and AMI[30,31]. Hence, we evaluated whether the atherosclerosis-perturbed network could be leveraged towards discovery of diagnostic blood biomarkers for CAD[32] and AMI[33] (**Fig. 1C;** *see Methods*). Altogether, 16 of the 31 genes in the atherosclerosis-perturbed network were also among the 6,346 differentially expressed genes (DEGs; adjusted p-value < 0.2) in transcriptomes of patients with CAD (**Fig. 1E**, GSE202625[32]) or AMI, (GSE66360[33]) (hypergeometric test p-value = 0.006). A disease score computed as the difference between the geometric mean of the 17 upregulated genes and the geometric mean of 14 downregulated genes[34] (see ***Extended Data Online Methods***) differentiated CAD (p-value = 0.0164) and AMI (p-value = 0.00057) patients from healthy controls in each respective cohort with area under the receiver operating characteristic curve (AUROC) > 0.7 (**Fig. 1F-G**). The likelihood of selecting a better performing 31-gene diagnostic signature for both CAD and AMI from the 5,041 genes transcriptionally profiled in both datasets and labeled as DEGs in at least one of the two datasets was 0.018. Notably, we observed similar or even better diagnostic performance with AUROCs ≥ 0.714 and ≥ 0.814, respectively, using each of three genes for CAD (ARAP2, P2RY14, and FKBP15) and two genes for AMI (SERPINA1, and ASGR2), from the atherosclerosis-perturbed network (**Fig. 1F-G**). The likelihood of finding better performing two and three genes from the 2,738 DEGs and 4,173 DEGs in the CAD and AMI datasets was ~0.003 and ~0.006, respectively. These findings were striking because the atherosclerosis-perturbed network was discovered with monocyte transcriptomes of patients stratified by CAC score, which by itself is an imperfect predictor of heart disease, with false negatives and false positives, due to challenges in score interpretation and variability with imaging techniques used[35,36]. Our findings demonstrate how monocyteMINER helped to decode transcriptome states captured within monocytes of a relatively small cohort of patients and healthy controls to uncover predictive blood biomarkers that were mechanistically associated with atherosclerosis, CAD, and AMI. In the next section, we describe how this monocyte network-based strategy was developed further and used to discover generalizable transcriptional biomarkers for active TB in whole blood.

#### Extended Data 3 | Benchmarking of Intermediate MAS_TB Signatures Against Prior Published Signatures

We benchmarked the performance of MAS_TB17, which was discovered using a cohort of just 11 patients, against four previously reported TB signatures (Sambarey16[37], Lee_4[38], Verhagen_10[39], and Leong_24[40]) that were also discovered using similarly small sample sizes (14-49 samples), across 14 cohorts that covered three disease contexts: aTB vs healthy/LTBI controls (10 cohorts with a total of 958 samples), aTB vs. ODs (5 cohorts with 1,469 samples), and in advanced HIV patients with and without aTB (2 cohorts of 56 samples; **Table S2**). MAS_TB17 demonstrated superior performance in differentiating aTB from OD, with significantly higher AUROC ($p \leq 0.05$ for ROC comparisons[41]) in 75% of comparisons (15/20) and was never outperformed across five validation cohorts. When distinguishing aTB from healthy/LTBI controls across 10 validation cohorts, MAS_TB17 performance was comparable ($p > 0.05$) in 52.5% of comparisons (21/40) and superior in 32.5% (13/40). Only two signatures (Leong_24[40] and Sambarey16[37]) outperformed MAS_TB17 in 15% of cases (6/40). Notably, while MAS_TB17 matched or exceeded other signatures for aTB vs healthy/LTBI discrimination, it excelled particularly in the aTB vs OD context –a condition for which it was not originally designed. Finally, MAS_TB17 outperformed all four signatures in identifying aTB cases within two cohorts of advanced HIV patients (**Table S2**). Together, these findings demonstrate the ability of monocyte-MINER to uncover sensitive and specific disease biomarkers using small datasets (just 11 patients, in this example).

To reduce the number of genes in the signature and facilitate clinical translation, we evaluated the performance of all doublet and triplet combinations of genes within MAS_TB17 on an independent compendium of whole blood transcriptomes from 54 patients with aTB diagnosis and 93 controls, including 69 patients with LTBI and 24 healthy individuals[42]. Through 1,000 iterations of random sampling, selecting 74 patients (i.e., 50% of the samples) in each iteration, while maintaining the original proportions of aTB and control cases, we computed AUROCs for all candidate signatures and identified two best performing three-gene signatures: MAS_TB3a (ANKRD22, AIM2, and VAMP5; average AUROC: 0.962, 95% percentile interval: 0.935-0.989) and MAS_TB3b (IGF2BP3, AIM2, and GBP5; average AUROC: 0.937, 95% percentile interval: 0.894-0.982) (*see* ***Methods;*** **Fig. 2A** and **Table 2**). Notably, ROC curves of MAS_TB3a and MAS_TB3b were statistically indistinguishable across seven cohorts, and both significantly outperformed MAS_TB17 (i.e., higher AUROC with Delong test[41] p-value $< 0.05$) in five and three out of the ten validation cohorts, respectively (**Fig. 2B** and **Table S3**).

Finally, we calculated weighted mean AUROC scores (wAUROC[43]) to compare the overall performance of the three MAS_TB signatures across the 10 validation cohorts, to account for differences in the numbers of patients in each cohort (**Fig. 2B** and **Table S3**). The wAUROCs of both MAS_TB3a (0.900) and MAS_TB3b (0.877) were greater than that of MAS_TB17 (0.849) in detecting aTB vs healthy/LTBI. Importantly, MAS_TB3a and MAS_TB3b achieved WHO TPP for three different types of TB screening tests in at least one validation cohort [44]. MAS_TB3a achieved minimal performance thresholds for the TPP for a high-sensitivity and high-specificity test (specificity ≥80% and ≥90% sensitivity), a high-sensitivity screening test (specificity ≥60% and ≥90% sensitivity), and a high-specificity screening test (specificity ≥98% and ≥60% sensitivity) in 30%, 50%, and 60% of the validation cohorts, respectively (**Data Set S2**). MAS_TB3b achieved the minimal TPPs in 10%, 80%, and 40% of the validation cohorts for the different screening tests (**Data Set S2**).

#### Extended Data 4 | Full Comparative Performance Assessment: MAS_TB_META5 vs. 34 Published Signatures

MAS_TB_META5, MAS_TB3a, MAS_TB_OD3, and eight of the other 34 previous signatures (Blankley_5[45], Long_RES_10[46], Natarajan_7[47], Rajan_HIV_5[48], Sambarey_HIV_10[37], Suliman_4[49], Thompson_9[50], and Zhao_NANO_6[51]) were classified as top performers in detecting aTB in at least 80% of all validation cohorts that included healthy and LTBI cases (**Fig. 3A**, and **Data Set S2**). MAS_TB_META5 and seven of the 34 signatures (Hoang_OD_3[52], LauxdaCosta_OD_3[53], Long_RES_10[46], Natarajan_7[47], Sambarey_HIV_10[37], Suliman_4[49], Suliman_RISK_2[49]) achieved the new WHO minimal TPP for a high-sensitivity screening test in seven of the validation cohorts (**Data Set S2**). While Zhao_NANO_6[51] ( together with MAS_TB3b and MAS_TB_OD3) achieved the minimal TPP in one additional aTB vs. healthy/LTBI cohort, it was derived from 12 TB signatures, including Sweeney _OD_3[54] and Kaforou_27[55], which were discovered using one of the validation cohorts (GSE37250) in this study. Thus, the performance assessment of Zhao_NANO_6 was confounded by lack of complete independence of the validation dataset from the training dataset. Yet, Zhao_NANO_6 was not among the top performers in the aTB vs OD context, in which it also did not achieve the WHO TPP (**Data Set S2**). In the context of aTB vs OD, MAS_TB_META5 was a top performing signature in two of three cohorts (GSE144127[52], GSE39940[56], and GSE37250[55]), achieving WHO minimal TPP for a high-sensitivity screening test in GSE144127 (**Data Set S2**). Only 4 of the 34 TB signatures (LauxDaCosta_OD3[53], Roe_3[57] and Kaul_3[58]) were among top performers across a handful more cohorts than MAS_TB_META5 in the aTB vs OD context. However, these four signatures had significant tradeoffs across the aTB vs healthy/LTBI contexts (**Fig. 3B, Fig. S2 and Data Set S2**), and with the exception of Gong_OD_4, they did not achieve the minimal WHO TPP for a prognostic test in more than one of the four TB progression validation cohorts. In comparison, MAS_TB_META5 achieved WHO TPP for a prognostic test in three out of the four validation cohorts (**Table S4**; in depth performance assessment of signatures in predicting TB progression is presented in the main text). Remarkably, 24 of the 34 TB signatures failed to achieve WHO TPP for a high-sensitivity screening test in any validation cohort for aTB vs OD (**Data Set S2**), underscoring the challenge in differentiating aTB from ODs, as compared to healthy/LTBI cases[59].

### Extended Data Online Methods | Atherosclerosis Network Discovery and CAD/AMI Validation Methods

##### Discovery of the atherosclerosis-perturbed monocyteMINER network

To evaluate the quality of monocyteMINER, we leveraged the scores for coronary artery calcium (CAC) scan, commonly used to estimate the risk of heart disease[4]. CAC scores were available for 1,141 out of 1,202 participants in the study and available in the metadata associated with compendium of monocyte transcriptomes[3]. To identify regulons (and their associated TFs) with differential activity that potentially drive the development of atherosclerosis, we compared the regulon eigengenes (a proxy for regulon activity) between patients with presumed high and low risk of heart disease (stratified by CAC scores). Eigengene of each regulon across all samples was calculated by MINER as the first component of a Principal Component Analysis of the expression matrix of member genes[1,2]. Risk stratification based on CAC score was performed per established guidelines[5]. Individuals with CAC scores = 0 were considered to be at the lowest risk for developing atherosclerosis, whereas individuals with CAC scores > 300 were considered at high risk. Regulons that were differentially active in patients with low and high risk of developing atherosclerosis were uncovered by comparisons of eigengenes of each regulon in the two groups stratified by CAC scores, as described above. Regulons with adjusted t-test p-values < 0.05 were considered differentially active. Atherosclerosis relevance of regulon member genes and their associated TFs was based on prior literature as described in the Results.

##### Assessment of clinical relevance of atherosclerosis-perturbed monocyte MINER network in the diagnosis of CAD and AMI.

Given the direct link between atherosclerosis and CAD and the reported association between CAC scores and risk of myocardial infarction[36], clinical relevance of the atherosclerosis-perturbed monocyte MINER network was evaluated using blood-derived transcriptomes from a CAD cohort[32] (GEO ID: GSE202625, n=52), and an acute myocardial infarction (AMI) cohort[33] (GEO ID: GSE66360, n = 99). Both cohorts included healthy controls without CAD or AMI. Donors of samples transcriptionally profiled in the GSE202625 dataset were classified by Andreini et al. in the healthy or CAD groups based on their coronary computed tomography angiography (CCTA) results and degree of coronary stenosis[32] (i.e., artery narrowing). CCTA detects both calcified and non-calcified lesions, offering a more accurate and comprehensive assessment of heart disease status than the CAC score[60,61]. Raw read counts (RNAseq) for 27 CAD cases and 25 controls blood transcriptomes were downloaded from GEO and analyzed using DESeq2[62] to perform differential expression analysis while correcting for multiple sequencing runs. A cutoff of adjusted p-value < 0.2 in the DESeq2 output was used to define DEGs. We used this relaxed cutoff for exploring overlap of the atherosclerosis-perturbed network and GSE202625 without discounting subtle changes in expression of genes associated with heart disease-perturbed subnetworks, as commonly done in pathway-level analysis[63]. A similar cutoff has been used in previous investigations of disease-related human transcriptomes[63], and as an alternative (with more control over the false discovery rate) to raw p-value cutoffs previously applied in the analysis of GSE2022625 and similar datasets[64,65]. The GSE66360 dataset included transcriptomes related to a more severe and potentially lethal manifestation of heart disease that requires immediate attention[33]. Specifically, GSE66360 consisted of transcriptional profiles of endothelial cells circulating in blood of 50 individuals with AMI and 49 control participants[33]. Pre-normalized microarray data from the original authors was used for differential expression analysis using a Bayesian T-test implemented in the Cyber-T tool[66]. DEGs were defined using an adjusted p-value < 0.2, as explained before. Given the central role of the interaction between endothelial cells and monocytes during the development of atherosclerosis[67,68], GSE66360 allowed us to evaluate the conservation of the heart disease driven transcriptional remodeling revealed by monocyteMINER across functionally related and interacting cell types. We used the hypergeometric test to calculate the significance of enrichment of 31 genes (out of the total 6,346 DEGs across AMI and CAD cohorts) within the atherosclerosis-perturbed network (**Fig. 1C**). Altogether, the 3,189 DEGs (identified with an adjusted p-value cutoff < 0.1) were marginally significant enriched within the atherosclerosis-perturbed network (hypergeometric test p-value = 0.073). The ability of the atherosclerosis-perturbed network to distinguish control individuals from individuals with CAD or AMI was evaluated using a score computed as the difference between the geometric mean of genes upregulated in the high-risk group (i.e., regulons # 869 and # 4143) and the geometric mean of downregulated genes in the same group (i.e., regulons # 3322 and # 5294), as previously done for other diseases[34]. For the GSE202625 dataset, we transformed the data into log2(counts per million +1) before computing the geometric means. For the GSE66360 dataset, we directly used the pre-normalized log-transformed microarray expression values for computing the disease score. We performed 10,000 iterations of random gene sampling to estimate the probability of selecting 31 genes from a total of 5,041 DEGs in AMI and CAD with better performance (i.e., ≥AUROC) than the 31 genes (17 upregulated and 14 downregulated) identified with monocyteMINER. We used a similar approach to estimate the probability of selecting at least three genes among 2,738 DEGs in CAD (GSE202625), and two genes out of 4,173 DEGs in AMI (GSE66360) with better performance (≥AUROC values of 0.794, and 0.888 for CAD and AMI, respectively) than the three- and two-gene signatures identified using monocyteMINER for diagnosis of the two diseases (**Fig 1F-G**).

#### REFERENCES


1. Wall, M. A. *et al.* Genetic program activity delineates risk, relapse, and therapy responsiveness in multiple myeloma. *npj Precision Oncology* **5**, 60 (2021).
2. Plaisier, C. L. *et al.* Causal mechanistic regulatory network for glioblastoma deciphered using systems genetics network analysis. *Cell systems* **3**, 172–186 (2016).
3. Liu, Y. *et al.* Blood monocyte transcriptome and epigenome analyses reveal loci associated with human atherosclerosis. *Nature communications* **8**, 393 (2017).
4. Gepner, A. D. *et al.* Comparison of carotid plaque score and coronary artery calcium score for predicting cardiovascular disease events: the multi-ethnic study of atherosclerosis. *Journal of the American Heart Association* **6**, e005179 (2017).
5. Detrano, R. *et al.* Coronary calcium as a predictor of coronary events in four racial or ethnic groups. *New England Journal of Medicine* **358**, 1336–1345 (2008).
6. Ruscica, M., Busnelli, M., Runfola, E., Corsini, A. & Sirtori, C. R. Impact of PPAR-alpha polymorphisms—the case of metabolic disorders and atherosclerosis. *International journal of molecular sciences* **20**, 4378 (2019).
7. Marx, N. *et al.* PPARα activators inhibit tissue factor expression and activity in human monocytes. *Circulation* **103**, 213–219 (2001).
8. Zandbergen, F. & Plutzky, J. PPARα in atherosclerosis and inflammation. *Biochimica et Biophysica Acta (BBA)-Molecular and Cell Biology of Lipids* **1771**, 972–982 (2007).
9. Zhao, Z.-W. *et al.* Heat shock protein 70 accelerates atherosclerosis by downregulating the expression of ABCA1 and ABCG1 through the JNK/Elk-1 pathway. *Biochimica et Biophysica Acta (BBA)-Molecular and Cell Biology of Lipids* **1863**, 806–822 (2018).
10. Yan, W., Li, D. & Zhou, X. Pravastatin attenuates the action of the ETS domain-containing protein ELK1 to prevent atherosclerosis in apolipoprotein E-knockout mice via modulation

of extracellular signal-regulated kinase 1/2 signal pathway. *Clinical and Experimental Pharmacology and Physiology* **44**, 344–352 (2017).

11. Sikorski, K., Wesoly, J. & Bluyssen, H. A. Data mining of atherosclerotic plaque transcriptomes predicts STAT1-dependent inflammatory signal integration in vascular disease. *International journal of molecular sciences* **15**, 14313–14331 (2014).
12. Eskandarian Boroujeni, M. *et al.* Integrative multi-omics analysis of IFNγ-induced macrophages and atherosclerotic plaques reveals macrophage-dependent STAT1-driven transcription in atherosclerosis. *Frontiers in Immunology* **16**, 1590953 (2025).
13. Liu, C. *et al.* Identifying RBM47, HCK, CD53, TYROBP, and HAVCR2 as hub genes in advanced atherosclerotic plaques by network-based analysis and validation. *Frontiers in Genetics* **11**, 602908 (2021).
14. Rommereim, L. M. *et al.* A small sustained increase in NOD1 abundance promotes ligand-independent inflammatory and oncogene transcriptional responses. *Science signaling* **13**, eaba3244 (2020).
15. Martín-Ventura, J. L. *et al.* Increased CD74 expression in human atherosclerotic plaques: contribution to inflammatory responses in vascular cells. *Cardiovascular research* **83**, 586–594 (2009).
16. Bernhagen, J. *et al.* MIF is a noncognate ligand of CXC chemokine receptors in inflammatory and atherogenic cell recruitment. *Nature medicine* **13**, 587–596 (2007).
17. Burger-Kentischer, A. *et al.* Expression of macrophage migration inhibitory factor in different stages of human atherosclerosis. *Circulation* **105**, 1561–1566 (2002).
18. Trifone, C. *et al.* Interaction between macrophage migration inhibitory factor and CD74 in human immunodeficiency virus type I infected primary monocyte-derived macrophages triggers the production of proinflammatory mediators and enhances infection of unactivated CD4+ T cells. *Frontiers in Immunology* **9**, 1494 (2018).

19. Metzler, B. *et al.* Activation of heat shock transcription factor 1 in atherosclerosis. *The American journal of pathology* **162**, 1669–1676 (2003).

20. Krishnamurthy, K. *et al.* Heat shock factor-1 knockout enhances cholesterol 7α-hydroxylase (CYP7A1) and multidrug transporter (MDR1) gene expressions to attenuate atherosclerosis. *Cardiovascular Research* **111**, 74–83 (2016).

21. Østerud, B. & Bjørklid, E. Role of monocytes in atherogenesis. *Physiological reviews* **83**, 1069–1112 (2003).

22. Gleissner, C. A., Leitinger, N. & Ley, K. Effects of native and modified low-density lipoproteins on monocyte recruitment in atherosclerosis. *Hypertension* **50**, 276–283 (2007).

23. Bekkering, S. *et al.* Oxidized low-density lipoprotein induces long-term proinflammatory cytokine production and foam cell formation via epigenetic reprogramming of monocytes. *Arteriosclerosis, thrombosis, and vascular biology* **34**, 1731–1738 (2014).

24. Frostegård, J. *et al.* Oxidized low density lipoprotein induces differentiation and adhesion of human monocytes and the monocytic cell line U937. *Proceedings of the National Academy of Sciences* **87**, 904–908 (1990).

25. Jego, G. *et al.* Dual regulation of SPI1/PU. 1 transcription factor by heat shock factor 1 (HSF1) during macrophage differentiation of monocytes. *Leukemia* **28**, 1676–1686 (2014).

26. Frostegård, J. *et al.* Induction of heat shock protein in monocytic cells by oxidized low density lipoprotein. *Atherosclerosis* **121**, 93–103 (1996).

27. Pockley, A. G. Heat shock proteins, inflammation, and cardiovascular disease. *Circulation* **105**, 1012–1017 (2002).

28. Hilgendorf, I., Swirski, F. K. & Robbins, C. S. Monocyte fate in atherosclerosis. *Arteriosclerosis, thrombosis, and vascular biology* **35**, 272–279 (2015).

29. Zhang, S.-M. *et al.* Interferon regulatory factor 9 is critical for neointima formation following vascular injury. *Nature communications* **5**, 5160 (2014).

30. Libby, P. & Theroux, P. Pathophysiology of coronary artery disease. *Circulation* **111**, 3481–3488 (2005).

31. Libby, P. The biology of atherosclerosis comes full circle: lessons for conquering cardiovascular disease. *Nature Reviews Cardiology* **18**, 683–684 (2021).

32. Andreini, D. *et al.* Whole-blood transcriptional profiles enable early prediction of the presence of coronary atherosclerosis and high-risk plaque features at coronary CT angiography. *Biomedicines* **10**, 1309 (2022).

33. Muse, E. D. *et al.* A whole blood molecular signature for acute myocardial infarction. *Scientific reports* **7**, 12268 (2017).

34. Chawla, D. G. *et al.* Benchmarking transcriptional host response signatures for infection diagnosis. *Cell Systems* **13**, 974–988 (2022).

35. Blaha, M. J., Mortensen, M. B., Kianoush, S., Tota-Maharaj, R. & Cainzos-Achirica, M. Coronary artery calcium scoring: is it time for a change in methodology? *JACC: Cardiovascular Imaging* **10**, 923–937 (2017).

36. Gupta, A. *et al.* Coronary artery calcium scoring: current status and future directions. *Radiographics* **42**, 947–967 (2022).

37. Sambarey, A. *et al.* Unbiased identification of blood-based biomarkers for pulmonary tuberculosis by modeling and mining molecular interaction networks. *EBioMedicine* **15**, 112–126 (2017).

38. Lee, S.-W. *et al.* Gene expression profiling identifies candidate biomarkers for active and latent tuberculosis. *BMC bioinformatics* **17**, S3 (2016).

39. Verhagen, L. M. *et al.* A predictive signature gene set for discriminating active from latent tuberculosis in Warao Amerindian children. *BMC genomics* **14**, 74 (2013).

40. Leong, S. *et al.* Existing blood transcriptional classifiers accurately discriminate active tuberculosis from latent infection in individuals from south India. *Tuberculosis* **109**, 41–51 (2018).

41. DeLong, E. R., DeLong, D. M. & Clarke-Pearson, D. L. Comparing the areas under two or more correlated receiver operating characteristic curves: a nonparametric approach. *Biometrics* 837–845 (1988).

42. Berry, M. P. *et al.* An interferon-inducible neutrophil-driven blood transcriptional signature in human tuberculosis. *Nature* **466**, 973–977 (2010).

43. Warsinske, H., Vashisht, R. & Khatri, P. Host-response-based gene signatures for tuberculosis diagnosis: A systematic comparison of 16 signatures. *PLoS medicine* **16**, e1002786 (2019).

44. World Health Organization. *Target Product Profile for Tuberculosis Screening Tests*. (World Health Organization, 2025).

45. Blankley, S. *et al.* A 380-gene meta-signature of active tuberculosis compared with healthy controls. *European Respiratory Journal* **47**, 1873–1876 (2016).

46. Long, N. P. *et al.* A 10-gene biosignature of tuberculosis treatment monitoring and treatment outcome prediction. *Tuberculosis* **131**, 102138 (2021).

47. Natarajan, S., Ranganathan, M., Hanna, L. E. & Tripathy, S. Transcriptional profiling and deriving a seven-gene signature that discriminates active and latent tuberculosis: an integrative bioinformatics approach. *Genes* **13**, 616 (2022).

48. Rajan, J. V. *et al.* A novel, 5-transcript, whole-blood gene-expression signature for tuberculosis screening among people living with human immunodeficiency virus. *Clinical Infectious Diseases* **69**, 77–83 (2019).

49. Suliman, S. *et al.* Four-gene pan-African blood signature predicts progression to tuberculosis. *American journal of respiratory and critical care medicine* **197**, 1198–1208 (2018).

50. Thompson, E. G. *et al.* Host blood RNA signatures predict the outcome of tuberculosis treatment. *Tuberculosis* **107**, 48–58 (2017).

51. Kaipilyawar, V. *et al.* Development and validation of a parsimonious tuberculosis gene signature using the digital NanoString nCounter platform. *Clinical Infectious Diseases* **75**, 1022–1030 (2022).

52. Hoang, L. T. *et al.* Transcriptomic signatures for diagnosing tuberculosis in clinical practice: a prospective, multicentre cohort study. *The Lancet Infectious Diseases* **21**, 366–375 (2021).

53. da Costa, L. L. *et al.* A real-time PCR signature to discriminate between tuberculosis and other pulmonary diseases. *Tuberculosis* **95**, 421–425 (2015).

54. Sweeney, T. E., Braviak, L., Tato, C. M. & Khatri, P. Genome-wide expression for diagnosis of pulmonary tuberculosis: a multicohort analysis. *The Lancet Respiratory Medicine* **4**, 213–224 (2016).

55. Kaforou, M. *et al.* Detection of tuberculosis in HIV-infected and-uninfected African adults using whole blood RNA expression signatures: a case-control study. *PLoS medicine* **10**, e1001538 (2013).

56. Anderson, S. T. *et al.* Diagnosis of childhood tuberculosis and host RNA expression in Africa. *New England Journal of Medicine* **370**, 1712–1723 (2014).

57. Roe, J. *et al.* Blood transcriptomic stratification of short-term risk in contacts of tuberculosis. *Clinical Infectious Diseases* **70**, 731–737 (2020).

58. Kaul, S. *et al.* Latent tuberculosis infection diagnosis among household contacts in a high tuberculosis-burden area: A comparison between transcript signature and interferon gamma release assay. *Microbiology Spectrum* **10**, e02445-21 (2022).

59. Drain, P. K. & Dalmat, R. R. The elusive allure of a rapid host blood signature for tuberculosis disease. *Journal of Clinical Microbiology* **62**, e01289-23 (2024).

60. Achenbach, S. & Raggi, P. Imaging of coronary atherosclerosis by computed tomography. *European heart journal* **31**, 1442–1448 (2010).

61. Eckert, J., Schmidt, M., Magedanz, A., Voigtländer, T. & Schmermund, A. Coronary CT angiography in managing atherosclerosis. *International journal of molecular sciences* **16**, 3740–3756 (2015).

62. Love, M. I., Huber, W. & Anders, S. Moderated estimation of fold change and dispersion for RNA-seq data with DESeq2. *Genome biology* **15**, 550 (2014).

63. Simmons, J. D. *et al.* Monocyte transcriptional responses to Mycobacterium tuberculosis associate with resistance to tuberculin skin test and interferon gamma release assay conversion. *Msphere* **7**, e00159-22 (2022).

64. Zhu, P. *et al.* Identification of 5 hub genes for diagnosis of coronary artery disease. *Frontiers in Cardiovascular Medicine* **10**, 1086127 (2023).

65. McCaffrey, T. A. *et al.* RNAseq profiling of blood from patients with coronary artery disease: signature of a T cell imbalance. *Journal of molecular and cellular cardiology plus* **4**, 100033 (2023).

66. Baldi, P. & Long, A. D. A Bayesian framework for the analysis of microarray expression data: regularized t-test and statistical inferences of gene changes. *Bioinformatics* **17**, 509–519 (2001).

67. Mestas, J. & Ley, K. Monocyte-endothelial cell interactions in the development of atherosclerosis. *Trends in cardiovascular medicine* **18**, 228–232 (2008).

68. Medrano-Bosch, M., Simón-Codina, B., Jiménez, W., Edelman, E. R. & Melgar-Lesmes, P. Monocyte-endothelial cell interactions in vascular and tissue remodeling. *Frontiers in Immunology* **14**, 1196033 (2023).
