## Supplementary Material for "Monocyte-amplified transcriptional signatures of human diseases"

### SUPPLEMENTAL FIGURES AND LEGENDS

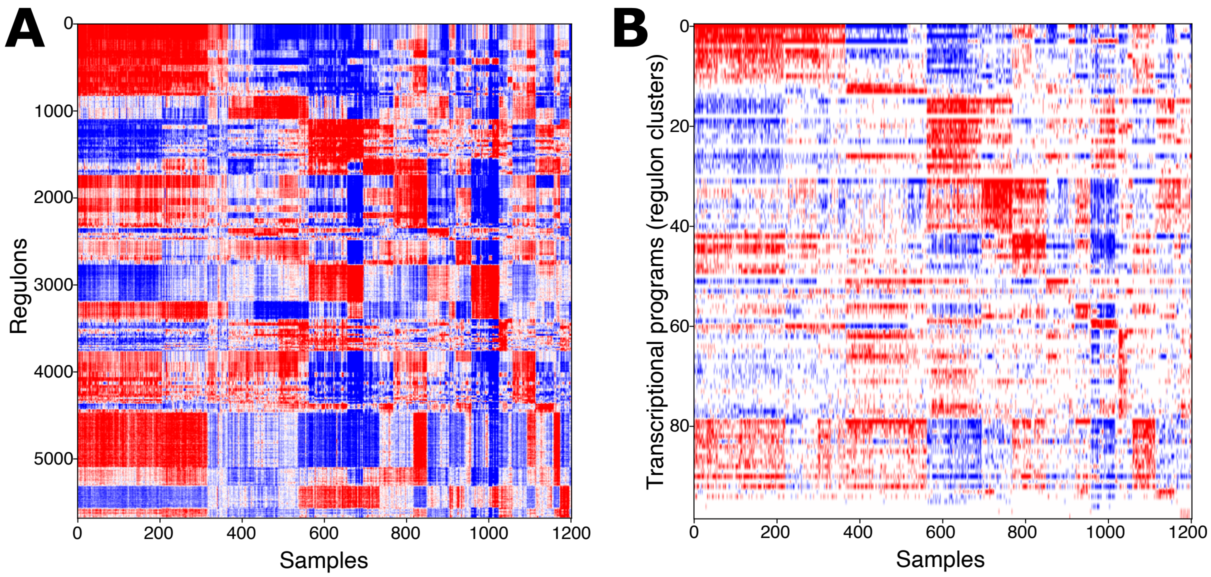


**Fig. S1. Activity of the monocyteMINER model across samples used for network inference.** A) Quantized activity of regulons (i.e., gene clusters) in monocyteMINER were classified as overactive (red), underactive (blue), or neutral (white) based on enrichment of each regulon with genes in the top third, bottom third, or intermediate range of distribution of z-scored expression[1] values across the MESA cohort[2]. B) Clustering of regulons with similar activity patterns across all patients identified 99 "*transcriptional programs*".

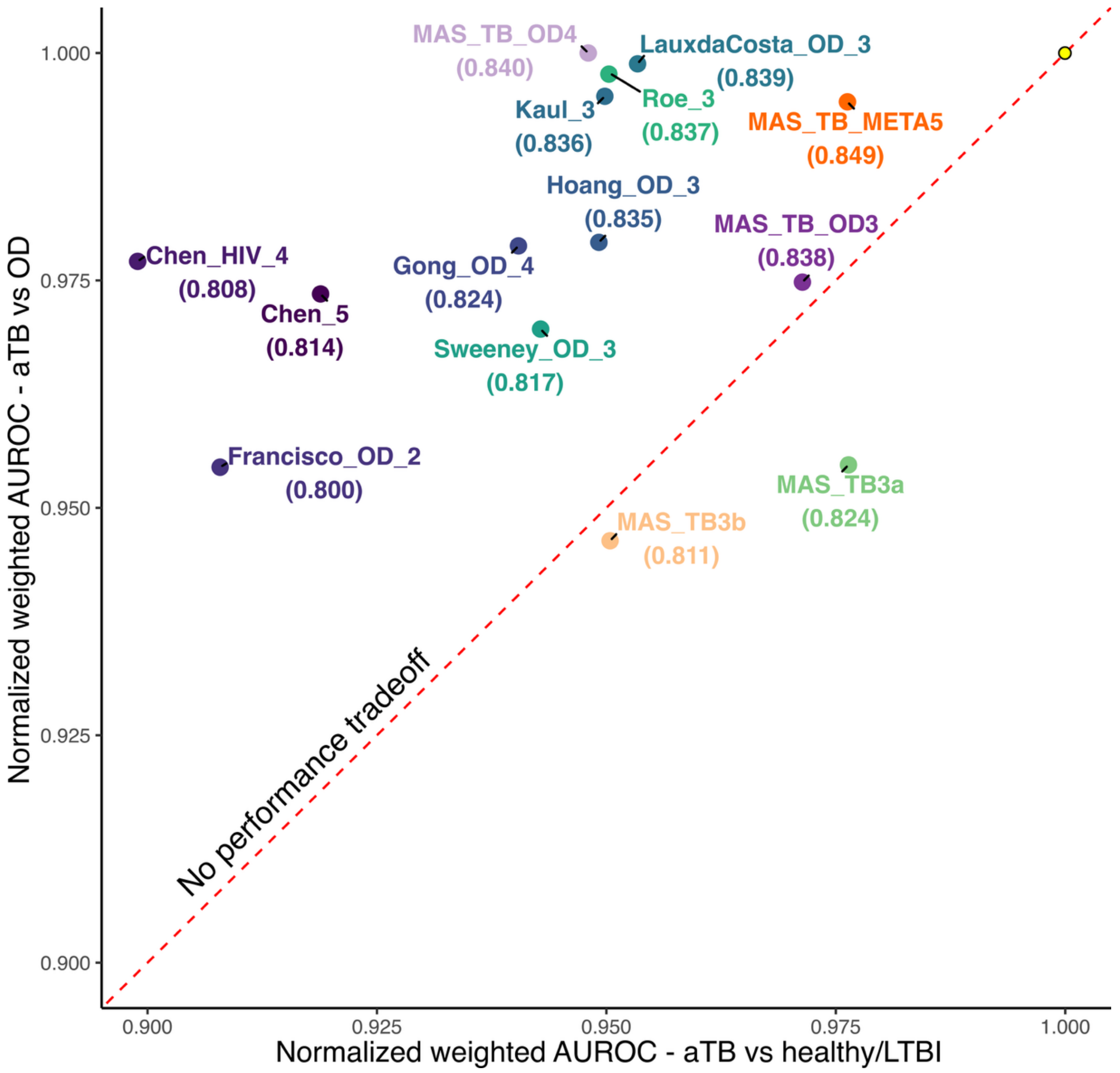


**Fig. S2. MAS_TB_META5 outperformed previous TB signatures at differentiating aTB from both healthy controls and OD even when considering TB cases without culture confirmation.** For each comparison (e.g., active TB vs other diseases), the overall performance of each signature across all validation cohorts was calculated as a weighted mean AUROC (wAUROC) score, as described in the main text, and normalized with respect to the maximum wAUROC across all TB signatures. Only signatures classified as top performing in more than one validation cohort for aTB vs OD (in addition to MAS_TB3a) are shown. The wAUROC scores in parentheses correspond to the overall performance of each signature across all validation cohorts for aTB vs healthy and aTB vs OD, after excluding datasets used for discovery or training. The dashed line represents the 1:1 line for equal performance in both disease contexts**.**

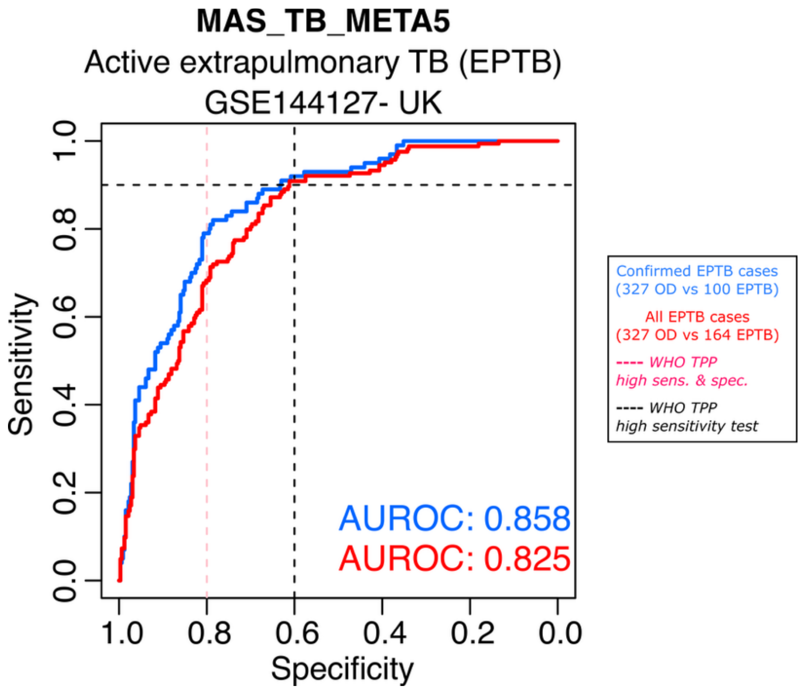


**Fig. S3. MAS_TB_META5 achieved WHO TPP for a high-sensitivity screening test for extrapulmonary TB (EPTB).** Performance of MAS_TB_META5 in detecting EPTB in a cohort that also included other non-TB diseases (OD) (GSE144127[3]) was evaluated using the PLAGE score[4] implementation in TBSignatureProfiler[5]. The red ROC curve shows diagnostic performance of MAS_TB_META5 when all EPTB cases were considered, and the blue ROC curve shows its diagnostic performance when only culture confirmed EPTB cases were included. In both cases, the performance of MAS_TB_META5 achieved WHO TPP for a high sensitivity screening test (top left quadrant delimited by the black dashed lines). Area under the ROC curve (AUROC) for each evaluation is shown in the bottom right corner.

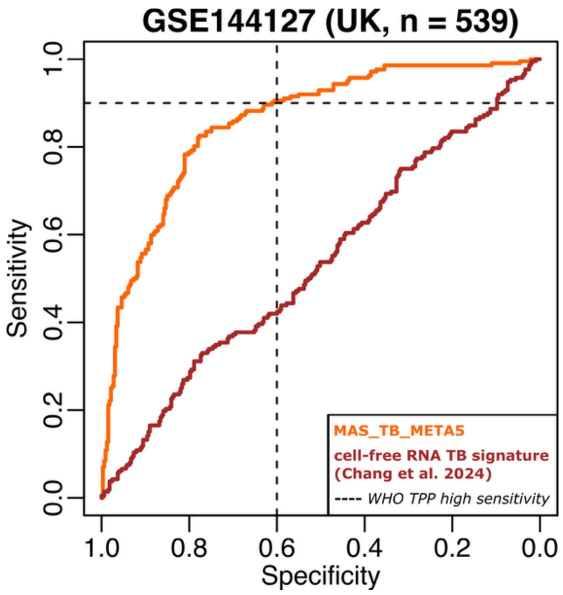


**Fig. S4. Comparative evaluation of MAS_TB_META5 and a 6-gene cell-free RNA TB signature[6] in detecting aTB vs. other diseases in whole blood.** Performance of the two signatures was evaluated using the PLAGE score[5] on the UK cohort of 212 confirmed TB cases and 327 cases of other diseases (GSE144127[3]). The 6-gene TB signature consists of GBP5, BNIP3L, KLF6, DYSF, LASP1, and PCBP1.

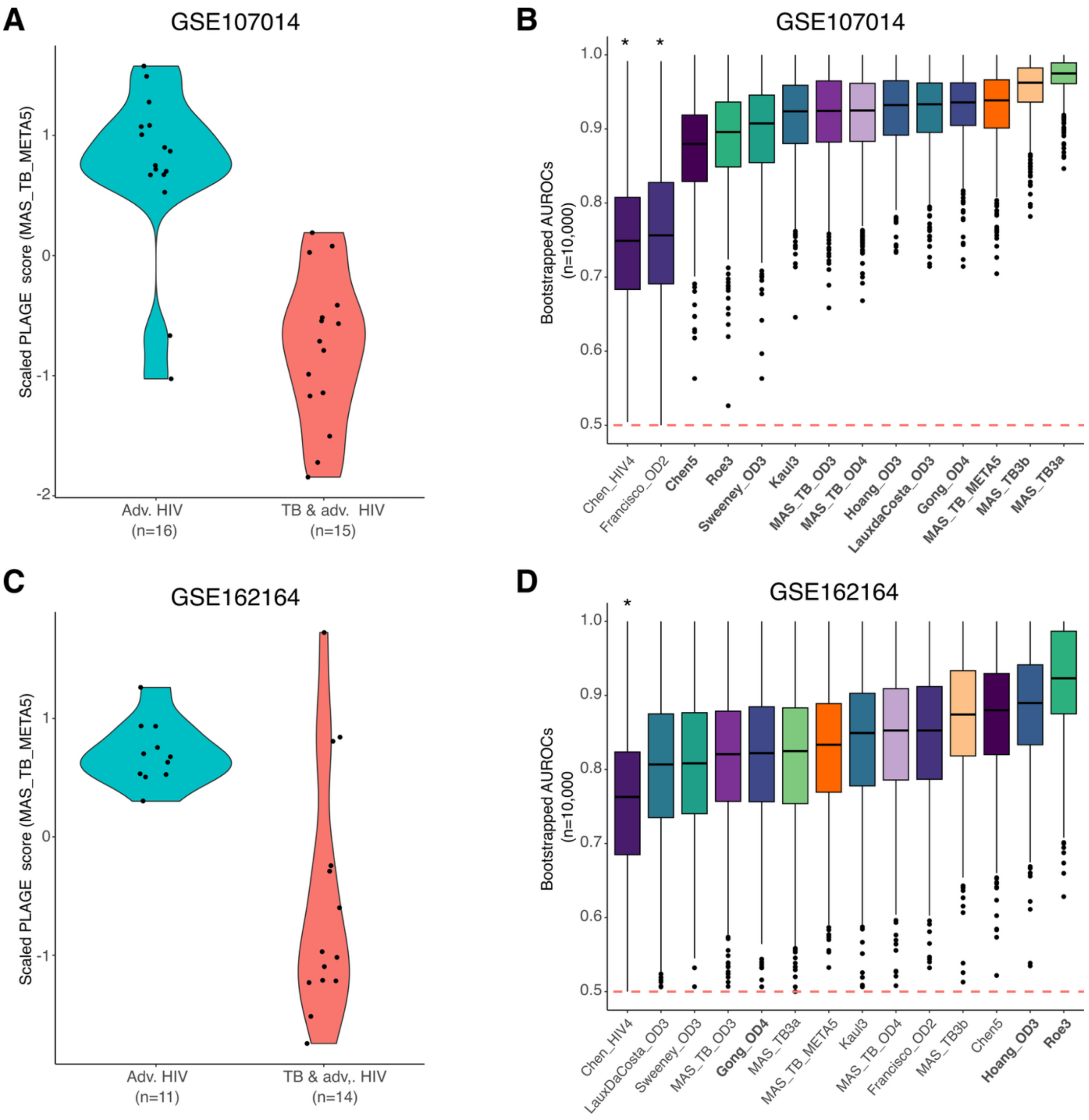


**E**

| Signature | AUROC[a] | | weighted AUROC |
|---|---|---|---|
| | GSE107014 | GSE162164 | |
| MAS_TB3b | 0.9542 (5.12e-08) | 0.8701 (0.0001) | 0.917 |
| Hoang_OD3 | 0.9208 (3.25e-06) | 0.8831 (0.0207) | 0.904 |
| MAS_TB3a | 0.9708 (1.02e-07) | 0.8182 (0.0005) | 0.903 |
| Roe3 | 0.8875 (4.92e-05) | 0.9156 (0.0104) | 0.900 |
| MAS_TB_META5 | 0.9292 (5.87e-07) | 0.8247 (0.0005) | 0.883 |
| MAS_TB_OD4 | 0.9167 (2.54e-05) | 0.8377 (0.0010) | 0.881 |
| Gong_OD4 | 0.9292 (2.56e-06) | 0.8182 (0.0612) | 0.880 |
| Kaul3 | 0.9125 (2.18e-06) | 0.8377 (0.0013) | 0.879 |
| Chen5 | 0.875 (0.001) | 0.8701 (0.0035) | 0.873 |
| LauxdaCosta_OD3 | 0.925 (1.95e-06) | 0.8052 (0.0009) | 0.872 |
| MAS_TB_OD3 | 0.9167 (6.71e-06) | 0.8117 (0.0007) | 0.870 |
| Sweeney_OD3 | 0.8958 (1.33e-05) | 0.8052 (0.0027) | 0.855 |
| Francisco_OD2 | 0.7583 (0.0139) | 0.8377 (0.0070) | 0.794 |
| Chen_HIV4 | 0.7458 (0.0204) | 0.7468 (0.0607) | 0.746 |

[a]TBSignatureProfiler P-values are shown in parentheses

**Fig. S5. Performance of 14 gene signatures, including MAS_TB_META5, in detecting aTB cases within two cohorts of advanced HIV patients (**GSE107104 and GSE162164**).** Violin plots of PLAGE scores for MAS_TB_META5 in a cohort of advanced 31 HIV patients from Uganda (GSE107104; **panel A**) and a cohort of 25 advanced HIV patients from India (GSE162164; **panel C**) with and without TB co-infection, before initiation of antiretroviral treatment. Performance of the 14 TB gene signatures in detecting aTB co-infections among 31 advanced HIV patients from Uganda (GSE107104; **panel B**) and 25 advanced HIV patients from India (GSE162164; **panel D**). Each boxplot indicates the distribution of AUROCs for a given signature across 10,000 bootstraps. Signatures were ranked by median AUROC, and shown in bold font if they achieved WHO TPP for a high-sensitivity screening test (sensitivity ≥ 90% and specificity ≥ 60%). Significant differences in AUROCs relative to best performing signature (highest median AUROC) are indicated with an asterisk (*). E) Individual and weighted mean AUROC values that underlie the boxplots for the 14 TB signatures in panels B and D.

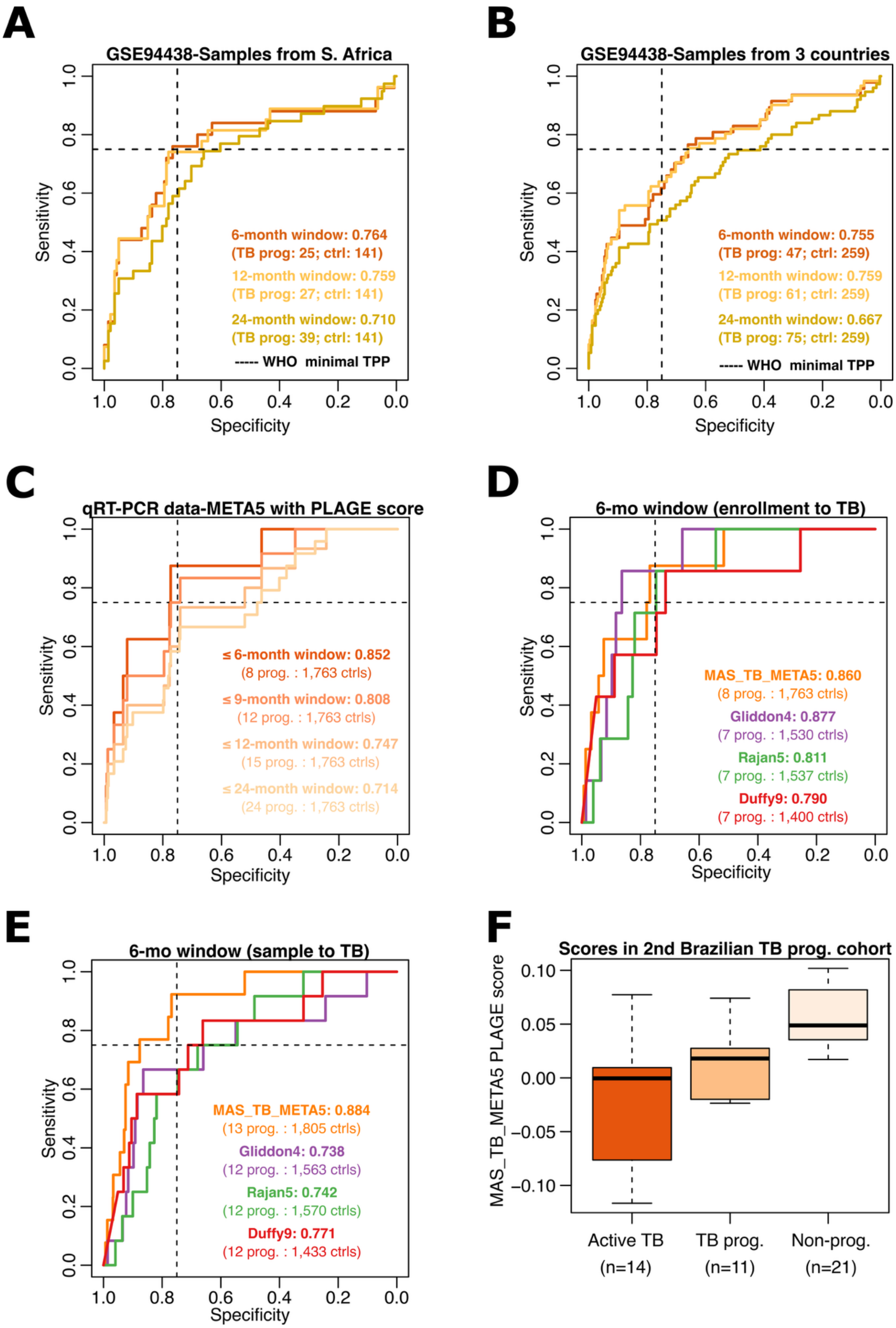
A
GSE94438-Samples from S. Africa
Sensitivity
Specificity
6-month window: 0.764
(TB prog: 25; ctrl: 141)
12-month window: 0.759
(TB prog: 27; ctrl: 141)
24-month window: 0.710
(TB prog: 39; ctrl: 141)
----- WHO minimal TPP
B
GSE94438-Samples from 3 countries
Sensitivity
Specificity
6-month window: 0.755
(TB prog: 47; ctrl: 259)
12-month window: 0.759
(TB prog: 61; ctrl: 259)
24-month window: 0.667
(TB prog: 75; ctrl: 259)
----- WHO minimal TPP
C
qRT-PCR data-META5 with PLAGE score
Sensitivity
Specificity
≤ 6-month window: 0.852
(8 prog. : 1,763 ctrls)
≤ 9-month window: 0.808
(12 prog. : 1,763 ctrls)
≤ 12-month window: 0.747
(15 prog. : 1,763 ctrls)
≤ 24-month window: 0.714
(24 prog. : 1,763 ctrls)
D
6-mo window (enrollment to TB)
Sensitivity
Specificity
MAS_TB_META5: 0.860
(8 prog. : 1,763 ctrls)
Gliddon4: 0.877
(7 prog. : 1,530 ctrls)
Rajan5: 0.811
(7 prog. : 1,537 ctrls)
Duffy9: 0.790
(7 prog. : 1,400 ctrls)
E
6-mo window (sample to TB)
Sensitivity
Specificity
MAS_TB_META5: 0.884
(13 prog. : 1,805 ctrls)
Gliddon4: 0.738
(12 prog. : 1,563 ctrls)
Rajan5: 0.742
(12 prog. : 1,570 ctrls)
Duffy9: 0.771
(12 prog. : 1,433 ctrls)
F
Scores in 2nd Brazilian TB prog. cohort
MAS_TB_META5 PLAGE score
Active TB
(n=14)
TB prog.
(n=11)
Non-prog.
(n=21)

**Fig. S6. Performance on MAS_TB_META5 in predicting TB progression at different time intervals in cohorts across diverse geographic locations.** A) Performance of MAS_TB_META5 (with PLAGE score) in predicting TB progression across a subset of 180 patients from S. Africa in the GSE94438 dataset at the 6-month, 12-month and 24-month intervals. B) Same as panel A across all 334 patients in the full GSE94438 dataset (from S. Africa, Ethiopia, and the Gambia). C) Performance of MAS_TB_META5 in predicting TB progression across 1,787 individuals (without missing Ct values for ANKRD22, VAMP5, and GBP5) from Brazil (Brazilian cohort A[7]) at the 6-month, 9-month, 12-month and 24-month intervals (the PLAGE score for each signature was calculated using relevant qRT-PCR Ct values). D) Performance of MAS_TB_META5 and three gene signatures (Gliddon4, Rajan5 and Duffy9) in predicting TB progression over a 6-month window post-enrollment in the Brazilian cohort A. While MAS_TB_META5 scores were computed as sum of delta Ct values of member genes, the normalized scores for Gliddon4, Rajan5 and Duffy9 used in this analysis were computed and published previously by Mendelsohn and colleagues[7]. E) Same as panel D but prognostic performance was assessed using pre-disease blood transcriptomes profiled up to 6-months prior to aTB diagnosis (see ***Methods***). F) Distribution of PLAGE scores for MAS_TB_META5 across 46 patients in the Brazilian cohort B (GSE112104). Number of individuals in each group are shown in parentheses. For visualization purposes, outliers are not shown. Note: Across panels A through E, the overall performance (i.e., AUROC) is indicated adjacent to the gene signature name or time interval. Numbers of progressors (prog.) and non-progressors (ctrl) are shown in parentheses.

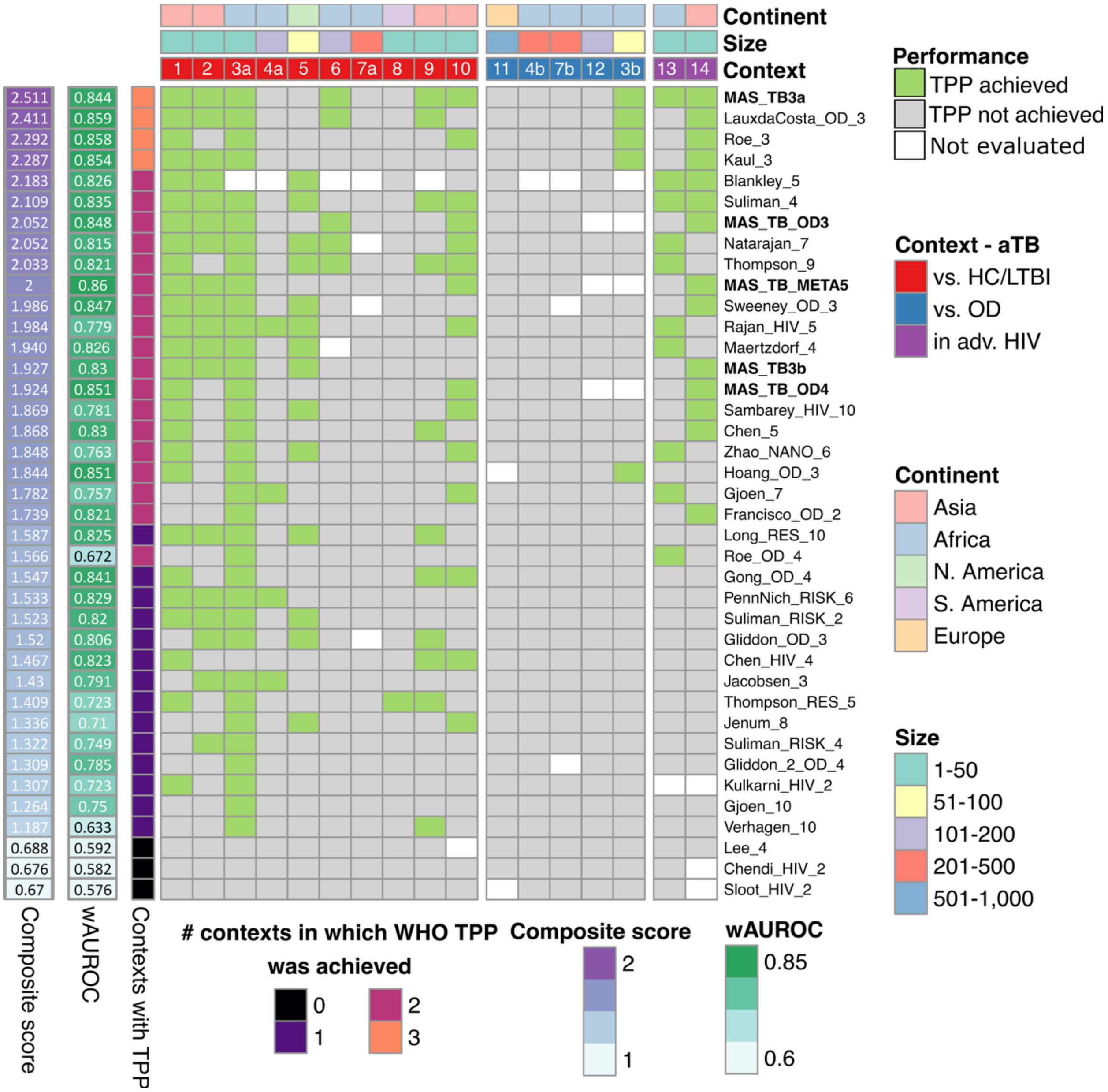


**Fig. S7. MAS_TB3a achieved WHO TPP for a high-specificity screening test across three different contexts.** Fourteen cohorts used for validation of signature performance are organized into three contexts (see "Context" track), with numbers of subjects in each cohort indicated in "Size" track, and geographic location noted in the "Continent" track. For datasets that contain aTB cases, OD cases, and healthy/latent TB infection cases, "a" indicates performance assessment in aTB vs. healthy/LTBI context, and "b" indicates assessment in aTB vs.OD context. Signatures are ranked based on a "Composite score", calculated as the sum of normalized values of "wAUROC" (to account for cohort size) and the number of cohorts and contexts in which WHO TPP was achieved

("Context with TPP"). MAS TB signatures are shown in bold font. In the heatmap, green shading indicates cohorts in which WHO TPP for a high-specificity screening test was achieved, gray shading indicates cohorts where WHO TPP was not achieved, and white indicates cohorts used for discovery of the relevant signature, and hence excluded from its performance assessment.

**SUPPLEMENTARY TABLES**

**Table S1- Functional enrichment of atherosclerosis-perturbed regulon # 869**

| Regulon | Functional term (GO Biological Process) | Enrichment adjusted p-value | Genes associated with enriched term |
|---|---|---|---|
| 869 | POSITIVE_REGULATION_OF_INTERLEUKIN_6_PRODUCTION | 0.0045 | NOD1, TYROBP, CD74 |
| 869 | POSITIVE_REGULATION_OF_DENDRITIC_CELL_ANTIGEN_PROCESSING_AND_PRESENTATION | 0.0045 | NOD1, CD74 |
| 869 | POSITIVE_REGULATION_OF_ANTIGEN_PROCESSING_AND_PRESENTATION | 0.0045 | NOD1, CD74 |
| 869 | DENDRITIC_CELL_ANTIGEN_PROCESSING_AND_PRESENTATION | 0.0045 | NOD1, CD74 |
| 869 | REGULATION_OF_DENDRITIC_CELL_ANTIGEN_PROCESSING_AND_PRESENTATION | 0.0045 | NOD1, CD74 |
| 869 | REGULATION_OF_ANTIGEN_PROCESSING_AND_PRESENTATION | 0.0122 | NOD1, CD74 |
| 869 | INTERLEUKIN_6_PRODUCTION | 0.0131 | NOD1, TYROBP, CD74 |
| 869 | POSITIVE_REGULATION_OF_HEMOPOIESIS | 0.0131 | TYROBP, CD74, PIK3R6 |
| 869 | POSITIVE_REGULATION_OF_MYELOID_LEUKOCYTE_DIFFERENTIATION | 0.0438 | TYROBP, CD74 |

Functional enrichment was evaluated using the ontology gene sets (C5) of the Human MSigDB collection[8].

**Table S2. Comparison of performance of MAS_TB17 and previous TB gene signatures discovered using small discovery cohorts.**

| Signature[a] | # samples used for discovery | Weighted AUROC[b,c] | | |
|---|---|---|---|---|
| | | aTB vs OD (N=1,558[d]) | aTB vs healthy/LTBI (N= 958[d]) | Advanced HIV-aTB co-infection vs HIV (N= 56[d]) |
| MAS_TB17 | 11 | 0.786 | 0.849 | 0.915 |
| Sambarey16[e] | 14 | 0.698 (2-0-3) | 0.893 (0-2-8) | 0.879 (0-0-2) |
| Lee_4 | 21 | 0.573 (5-0-0) | 0.615 (8-0-2*) | 0.719 (1-0-1) |
| Verhagen_10 | 27 | 0.558 (5-0-0) | 0.741 (4-0-6) | 0.740 (1-0-1) |
| Leong_24 | 44 | 0.693 (3-0-2) | 0.910 (1-4-5*) | 0.884 (0-0-2) |

[a]The number of genes is shown at the end of the name of each signature
[b]Weighted AUROC was computed as proposed by Warsinske et al.[9]
[c]In parenthesis is the number of validation cohorts in which the AUROC of previously proposed TB signatures was significantly lower than, higher than or not significantly different than the AUROC of MAS_TB17, respectively.
[d]The total number of samples (N) across the validation cohorts in each context is indicated in parentheses.
[e]Not included among TB signatures that were pre-compiled in the TBSignatureProfiler. The signature was specifically added for this analysis.
*This group includes a cohort used for the discovery of the TB signature being compared with MAS_TB17

**Table S3. Comparison of the AUROC of MAS_TB17, MAS_TB3a, MAS_TB3b and MAS_TB_META5 across 10 aTB vs healthy/LTBI validation cohorts[a]**

| Validation cohort (country) | MAS_TB17 vs MAS_TB3a | MAS_TB17 vs MAS_TB3b | MAS_TB3a vs MAS_TB3b | MAS_TB3a vs MAS_TB_META5 |
|---|---|---|---|---|
| GSE101705 (India) | N.S. | N.S. | N.S. | N.S. |
| GSE152218 (India) | MAS_TB3a | N.S. | N.S. | N.S. |
| GSE39939 (Kenya) | N.S. | N.S. | N.S. | N.S. |
| GSE39940 (S. Africa & Malawi) | N.S. | MAS_TB3b | MAS_TB3b | N.S. |
| GSE73408 (USA) | MAS_TB3a | N.S. | N.S. | N.S. |
| GSE28623 (Gambia) | MAS_TB3a | MAS_TB3b | N.S. | MAS_TB_META5 |
| GSE37250 (S. Africa & Malawi) | MAS_TB3a | MAS_TB3b | MAS_TB3a | N.S. |
| GSE84076 (Brazil) | N.S. | N.S. | N.S. | N.S. |
| GSE56153 (Indonesia) | N.S. | MAS_TB17 | N.S. | N.S. |
| GSE62525 (Taiwan) | MAS_TB3a | N.S. | MAS_TB3a | N.S. |
| [a]For each pair-wise ROC comparison, the signature with significantly higher AUROC is listed. Instances of no significant difference (i.e., p-value >0.05) are indicated as "N.S.". | | | | |

**Table S4: Comparison of MAS_TB_META5 and selected previous signatures for predicting risk of TB progression across four different cohorts[a,b]**

| Signature | wAUROC[c] | GSE94438[d] (6 months) | Mendelsohn[e] (9 months) | GSE79362 (2 years) | GSE112104 (5 years) |
|---|---|---|---|---|---|
| MAS_TB_META5 | 0.852 | Yes | Yes | No | Yes |
| LauxDaCosta_OD3 | 0.845 | No | No | No | No |
| Gong_OD4 | 0.837 | Yes | Yes[f] | No | Yes |
| Roe_3 | 0.849 | No | No | NA[g] | No |

| | | | | | |
|---|---|---|---|---|---|
| Kaul_3 | 0.843 | No | Yes | No | No |

[a]WHO target product profile (TPP) for a prognostic test (≥ 75% sensitivity and ≥ 75% specificity) was used as the performance metric. Instances in which a gene signature achieved WHO TPP when evaluating performance with the PLAGE score are indicated as “Yes”, while cases where WHO TPP were not achieved are annotated as “No”.

[b]For each cohort, time window for confirmed TB progression events are indicated in parentheses.

[c]Weighted AUROC was computed as proposed by Warsinske et al.[9] combining performance in the aTB vs healthy/LTBI, aTB vs OD contexts, TB and advanced HIV co-infection context, and the TB progression context. All datasets except Mendelsohn et al. (a qRT-PCR dataset for the TB progression context) were used to compute the wAUROCs. For each signature, only validation datasets that included at least two genes of the signatures and were not used for signature discovery or training were included in the wAUROC calculation.

[d]Analysis restricted to the South African subgroup, the largest of the three in the Pan-African dataset.

[e]For previous TB signatures, unless indicated otherwise, normalized scores computed and published by Mendelsohn et al. were used to evaluate performance. For MAS_TB_META5 and Kaul_3, each sample was scored as the sum of delta Ct values for ANKRD22, VAMP5 and GBP5, and FCGR1B, GBP1 and GBP5, respectively.

[f]Disease score was computed using geometric mean (as proposed by Gong et al.[10]) of delta Ct of BATF2, VAMP5, and SERPING1. UBE2L6, the other gene in Gong_OD4, was not present in the Mendelsohn dataset.

[g]The Roe_3 signature was discovered using the GSE79362 dataset. Thus, GSE79362 was not used as a validation cohort for Roe_3.

### REFERENCES


1. Wall, M. A. *et al.* Genetic program activity delineates risk, relapse, and therapy responsiveness in multiple myeloma. *npj Precision Oncology* **5**, 60 (2021).
2. Liu, Y. *et al.* Blood monocyte transcriptome and epigenome analyses reveal loci associated with human atherosclerosis. *Nature communications* **8**, 393 (2017).
3. Hoang, L. T. *et al.* Transcriptomic signatures for diagnosing tuberculosis in clinical practice: a prospective, multicentre cohort study. *The Lancet Infectious Diseases* **21**, 366–375 (2021).
4. Tomfohr, J., Lu, J. & Kepler, T. B. Pathway level analysis of gene expression using singular value decomposition. *BMC bioinformatics* **6**, 1–11 (2005).
5. Johnson, W. E. *et al.* Comparing tuberculosis gene signatures in malnourished individuals using the TBSignatureProfiler. *BMC infectious diseases* **21**, 1–13 (2021).
6. Chang, A. *et al.* Circulating cell-free RNA in blood as a host response biomarker for detection of tuberculosis. *Nature Communications* **15**, 4949 (2024).
7. Mendelsohn, S. C. *et al.* Transcriptomic Signatures of Progression to Tuberculosis Disease Among Close Contacts in Brazil. *The Journal of Infectious Diseases* jiae237 (2024).
8. Liberzon, A. *et al.* Molecular signatures database (MSigDB) 3.0. *Bioinformatics* **27**, 1739–1740 (2011).
9. Warsinske, H., Vashisht, R. & Khatri, P. Host-response-based gene signatures for tuberculosis diagnosis: A systematic comparison of 16 signatures. *PLoS medicine* **16**, e1002786 (2019).
10. Gong, Z. *et al.* The evaluation and validation of blood-derived novel biomarkers for precise and rapid diagnosis of tuberculosis in areas with high-TB burden. *Frontiers in Microbiology* **12**, 650567 (2021).
